# The longitudinal dynamics and natural history of clonal haematopoiesis

**DOI:** 10.1101/2021.08.12.455048

**Authors:** Margarete A. Fabre, José Guilherme de Almeida, Edoardo Fiorillo, Emily Mitchell, Aristi Damaskou, Justyna Rak, Valeria Orrù, Michele Marongiu, MS Vijayabaskar, Joanna Baxter, Claire Hardy, Federico Abascal, Michael Spencer Chapman, Nicholas Williams, Jyoti Nangalia, Iñigo Martincorena, Peter J. Campbell, Eoin F. McKinney, Francesco Cucca, Moritz Gerstung, George S. Vassiliou.

## Abstract

Human cells acquire somatic mutations throughout life, some of which can drive clonal expansion. Such expansions are frequent in the haematopoietic system of healthy individuals and have been termed clonal haematopoiesis (CH). While CH predisposes to myeloid neoplasia and other diseases, we have limited understanding of how and when CH develops, what factors govern its behaviour, how it interacts with ageing and how these variables relate to malignant progression. Here, we track 697 CH clones from 385 individuals aged 55 or older over a median of 13 years. We find that 92.4% of clones expanded at a stable exponential rate over the study period, with different mutations driving substantially different growth rates, ranging from 5% (*DNMT3A*, *TP53*) to over 50%/yr (*SRSF2-*P95H). Growth rates of clones with the same mutation differed by approximately +/−5%/yr, proportionately impacting “slow” drivers more substantially. By combining our time-series data with phylogenetic analysis of 1,731 whole genome-sequenced haematopoietic colonies from 7 older individuals, we reveal distinct patterns of lifelong clonal behaviour. *DNMT3A*-mutant clones preferentially expanded early in life and displayed slower growth in old age, in the context of an increasingly competitive oligoclonal landscape. By contrast, splicing gene mutations only drove expansion later in life, while growth of *TET2*-mutant clones showed minimal age-dependency. Finally, we show that mutations driving faster clonal growth carry a higher risk of malignant progression. Our findings characterise the lifelong natural history of CH and give fundamental insights into the interactions between somatic mutation, ageing and clonal selection.

## Introduction

Human haematopoiesis produces hundreds of billions of specialized blood cells every day, through a hierarchy of progressively more differentiated and numerous cells originating from a pool of long-lived haematopoietic stem cells (HSCs). Haematopoiesis remains highly efficient for decades, but is inevitably challenged by the phthisic effects of ageing^1–3^ and the inexorable acquisition of somatic DNA mutations^4^. Mutations that augment HSC “fitness” can drive clonal expansion of a mutant HSC and its progeny, a phenomenon known as clonal haematopoiesis (CH)^5–8^. CH becomes ubiquitous with advancing age and is associated with an increased risk of myeloid leukaemias and some non-haematological diseases^5–7,9–11^.

The observation that CH-associated mutations affect a very restricted set of genes that are also frequently mutated in leukaemia - most commonly those involved in epigenetic regulation (*DNMT3A*, *TET2* and *ASXL1*), splicing (*SF3B1* and *SRSF2*) and apoptosis (*TP53* and *PPM1D*)^5–8^ - implies that these mutations inherently confer fitness to HSCs. In fact, recent evolutionary models propose that each specific mutation carries a fixed fitness advantage, and that this explains the relative proportions and clonal sizes of CH driven by different driver mutations^12^. However, several observations suggest that non-mutation factors are also influential. For example, a handful of CH cases studied at two time-points propose that the clones driven by the same or very similar mutations can behave differently between individuals^11,13^. Also, the relative prevalence of different CH-driver gene mutations changes significantly depending on context; for example, in aplastic anaemia CH is commonly driven by mutations that enhance immune evasion^14–17^, whereas genotoxic stress favours clones with mutations in DNA damage genes^18–20^. Furthermore, factors like inflammation^21^ and heritable genetic variation^22–24^ can affect CH emergence.

A major limitation to our understanding of the determinants of CH behaviour/fate to date has been its reliance on cross-sectional studies capturing CH at single time-points. Here, by tracking blood cell clones over long periods of time in a large cohort, and by reconstructing haematopoietic phylogenies, we uncover the lifelong dynamics and natural history of CH.

## Results

### The age-dependent mutational landscape of CH

We analysed 1,593 blood DNA samples from 385 adults aged 54-93 years at the time of entry into the SardiNIA longitudinal study^25^. The participants, who had no history of haematological malignancy, were sampled up to 5 times (median 4) over 3.2-16 years (median 12.9 years) (Figure 1a, Extended Data Figure 1a-c). We performed deep targeted sequencing (mean 1,065x) of 56 genes associated with CH and haematological malignancy (Supplementary Table 1) and identified somatic mutations in 52 genes (Supplementary Table 2). Using the dNdScv algorithm, an implementation of dN/dS that corrects for trinucleotide mutation rates, sequence composition, and variable mutation rates across genes, we identified positive selection of missense and/or truncating variants in 17 of these genes (dN/dS ratio>1 with q<0.1) (Supplementary Table 3, Extended Data Figs. 2,3)^26^. We focussed on these genes for further analysis.

**Fig. 1:**
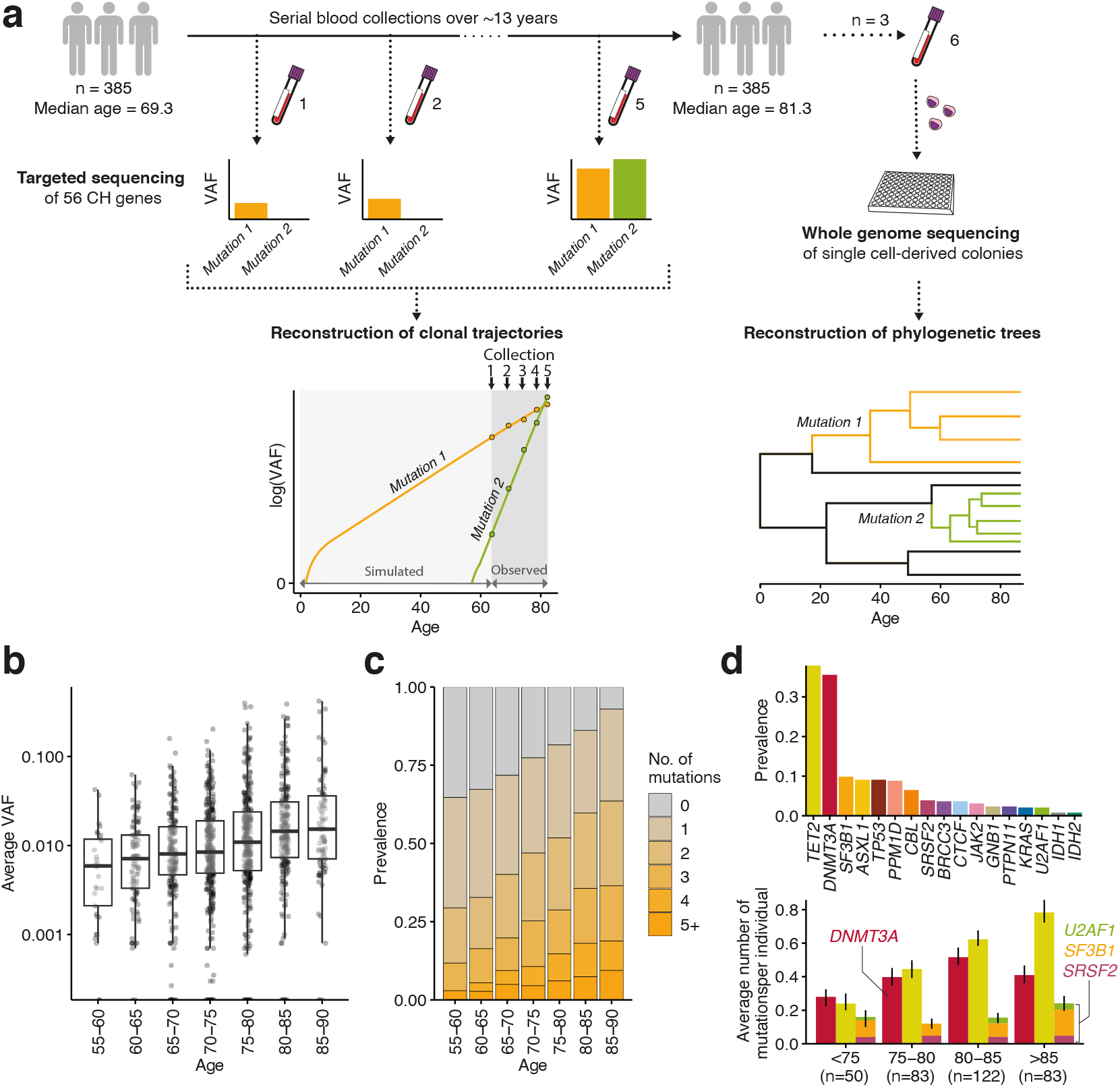
Experimental workflow and CH mutation characteristics. **a**, Study outline: 1,593 blood DNA samples were obtained from 385 elderly individuals sampled 2-5 times (median 4) over 3.2-16 years (median 12.9) and sequenced for mutations in 56 CH genes. Measured variant allele fractions (VAFs) were used to (i) fit observed clonal trajectories, and (ii) extrapolate the clonal dynamics prior to the period of observation. Additional blood samples from 3 selected individuals were used to generate 288 (3×96) whole-genome sequenced single cell-derived colonies for phylogeny reconstructions. **b**, Age distribution of average VAF per individual. **c,** Age-stratified prevalence of the number of mutations per individual. **d,** Prevalence of mutations in driver genes: upper panel shows absolute prevalence in the cohort; lower panel shows average number of mutations per individual in *DNMT3A*, *TET2* and splicing genes (*SF3B1, SRSF2, U2AF1*) at different ages.

At least one somatic non-synonymous mutation was identified in 305 of 385 individuals (79.2%), with CH prevalence, average clone size and number of mutations per individual increasing with advancing age, and CH identified in >90% of those aged 85 years or older (Fig. 1b,c). Mutations were most common in epigenetic regulator genes *TET2* and *DNMT3A*, and also frequent in *ASXL1*, *TP53*, *PPM1D* and spliceosome genes (Fig. 1d, upper panel). Interestingly, in this elderly cohort, advancing age impacted the prevalence of different driver mutations in a gene-dependent manner (Fig. 1d, lower panel). In particular, the prevalence of *DNMT3A* mutations showed no significant relationship with age overall (p=0.12, binomial regression of prevalence vs age, controlling for sex), whilst *TET2* mutations showed a consistent rise with age, averaging at 6.8%/yr (p=0.00037), as did mutations in splicing genes (*U2AF1*, *SRSF2* and *SF3B1*), whose prevalence increased by 5.4%/yr (p=0.025).

### Most clones expand steadily during older age

To investigate clonal behaviour over time, we used serial Variant Allele Fraction (VAF; the fraction of sequencing reads reporting a mutation) measurements as a surrogate for clone size, and fitted a saturating (logistic) exponential curve with a constant growth rate over time to each clonal trajectory. Such logistic growth behaviour is supported by simulations of evolutionary dynamics using Wright-Fisher models with constant fitness (Extended Data Fig. 4a-b)^27^. Remarkably, by assessing the fit between serial VAF measurements and the trajectories inferred by our model, we find that the great majority of clones (92.4%) expanded at a constant exponential rate over the study period (Fig. 2a,b, Extended Data Fig. 4c). The predominance of fixed-rate growth was particularly striking for genes like *DNMT3A* and *TET2*, for which 99% and 94.3% of clones, respectively, grew steadily over time. Nevertheless, some clones behaved unpredictably, with proportions varying by mutant gene. Most notable were *JAK2*-V617F-mutant clones, for which growth trajectories were particularly erratic, with only 58% displaying stable growth. The likelihood of mutant clones displaying non-constant growth at older age was not related to the number of mutations in the same individual (p=0.68; Extended Data Fig. 4d).

**Fig. 2:**
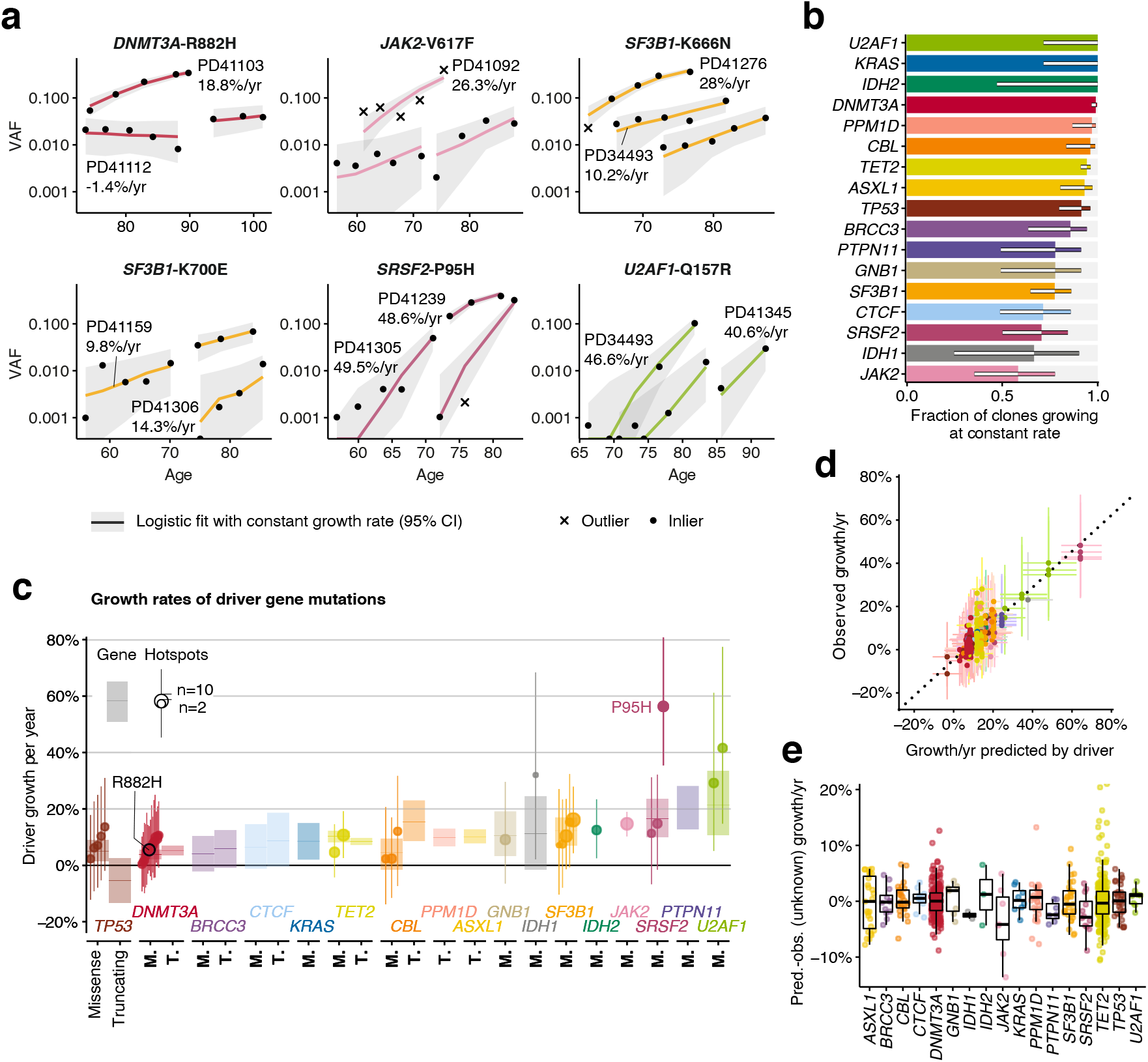
The longitudinal dynamics of CH in older age. **a,** Examples of fitted exponential growth of clones with mutations at 6 common hotspots. Grey bands represent the 95% highest posterior density interval (HPDI). Each data point is represented by a dot if it conforms to our model of fixed-rate exponential growth and by a cross otherwise (outlier, defined as tail probability less than 2.5%). **b,** Proportion of clonal trajectories showing fixed-rate exponential growth, ie. those with no outlying data-points as defined in (a), with 90% confidence intervals. **c,** Annual clonal growth associated with different driver mutations, for both whole genes and specific mutation sites. For gene-wise growth, truncating and missense mutations are modelled separately for genes where both are enriched. Sites are modelled separately to gene if mutated recurrently within our cohort. Point estimates for growth and 90% HDPI are represented for each site (dot and line, respectively, with dot size proportional recurrence) and each gene (horizontal line and rectangle, respectively). **d**, Relationship between clonal growth predicted by the identity of the driver mutation and actual observed growth, with 90% HDPI represented by vertical and horizontal lines, respectively. Vertical spread thereby captures differences in growth rate between clones bearing the same driver mutation. Clones growing faster than predicted lie above the dashed line, and slower clones lie below. **e**, Distribution of the unknown-cause effect for different genes. Each point represents a single clone and boxplots represent the distribution of these effects for each gene. The value of unknown-cause growth is *positive* for clones growing faster than expected by the identity of the driver mutation, and *negative* for clones growing slower than expected.

We further assessed the consistency of clonal trajectories by testing our ability to predict future clonal growth. Using additional prospectively-obtained blood samples from 11 individuals, we compared observed versus predicted VAFs (Extended Data Fig. 4e-g, Supplementary Table 4) and found good concordance (mean absolute error: 3.5%), corroborating our model and providing further evidence that fixed-rate growth is the norm in old age.

### Determinants of clonal growth rate

To delineate the factors that determine each clone’s growth rate, our logistic regression model fits the following contributions of the driver mutation: i) mutated gene; ii) specific amino acid change (for recurrently mutated sites) and iii) mutation type (truncating versus non-truncating) (Supplementary Table 5). An additional component in our model, measuring variation not captured by (i-iii), was also used and termed “unknown-cause growth” (Extended Data Fig. 4h).

We found that clones bearing mutations in different genes expanded at different rates, with mutations affecting *DNMT3A* and *TP53* displaying the slowest average annual growth rates of ~5% (Fig. 2c, Supplementary Table 6). Clones with mutations in the other most common driver genes (*TET2*, *ASXL1*, *PPM1D* and *SF3B1*), expanded at roughly twice this rate, i.e. ~10%/yr. The most rapidly expanding clones were those carrying mutations in *SRSF2*, *PTPN11* and *U2AF1*, growing at over 15-20%/yr on average. The only specific mutation displaying distinctive behaviour was *SRSF2*-P95H, which was associated with significantly faster expansion compared to other *SRSF2* mutations. By contrast, all other hotspot mutations drove growth at rates similar to mutations elsewhere in the same gene, including commonly mutated sites such as *DNMT3A*-R882, *SF3B1-*K666N and *SF3B1*-K700E.

For most genes, truncating and missense mutations drove comparable rates of growth. Exceptions were TP53, where missense grew 10%/yr (90% CI=[3-18%]) faster than truncating mutations (which usually did not expand or even contracted) and CBL, where missense grew 11%/yr (90% CI=[3-19%]) slower than truncating mutations (Fig. 2c, Extended Data Fig. 4i, Supplementary Table 6).

To quantify the impact of factors other than driver mutations, we compared the observed growth rate of each clone with that predicted by the mutation (Fig. 2d). In Figure 2d, vertical spread thereby represents the variability in growth rate between distinct clones with the same driver mutation. On average, this unknown-cause growth contributed approximately +/− 5%/yr to clonal expansion (Fig. 2e). Consequently, for fast-growing clones, including those associated with *SRSF2-*P95H or mutant *U2AF1*, this effect was proportionately small and there was relatively little inter-individual variability in growth rate. By contrast, the impact on “slow” drivers, such as *DNMT3A*, was more substantial, with some clones growing twice as fast as predicted by the mutation, and others showing negligible expansion. Clones harbouring *JAK2*-V617F mutations were an exception as they displayed an unusually high degree of inter-individual variability in relation to average growth rate (Fig. 2d,e, Extended Data Fig. 5a). In view of the well-described heritable contribution to myeloproliferative neoplasm (MPN) susceptibility^22,23^, we tested if *JAK2*-V617F-mutant clones grew faster in individuals with inherited MPN risk alleles, but found no such relationship (Extended Data Fig. 5b, Supplementary Table 7). However, we also made the more general observation that certain individuals harboured more mutations in the same gene than would be expected by chance (Extended Data Fig. 5c), suggesting that non-mutation factors influencing clonal growth are both individual- and gene-specific. While we found no evidence that these non-mutation factors include either sex or smoking history, since neither accounted for differences in clonal growth rate between individuals with the same mutant driver gene, age was a significant factor specifically for *TET2*-mutant clones, which grew faster in older individuals (Spearman’s rho=0.31; adj. p-value=2.33*10^−6^) (Extended Data Fig. 5d-f).

### Haematopoietic phylogenies give insights into the lifelong natural history of CH

To contrast the longitudinal clonal behaviours we observed in older age with lifelong clonal dynamics, we began by deriving and whole-genome sequencing (WGS) 96 single-cell-derived colonies from each of three individuals with splicing gene mutations (Fig. 3a-c), particularly as previous reports suggested a possible interaction of these mutations with age^8^. We constructed phylogenetic trees using somatic mutations as lineage-tracing barcodes and, since HSCs accumulate mutations at a near constant rate, we used phylogenetic branch lengths to time the onset of clonal expansions (“clades”)^28–31^. In PD41276, the phylogeny was dominated by an *SF3B1*-K666N-mutant clone, beginning between 23-47 years of age, with only a single *SF3B1*-wild type colony, consistent with a near-complete clonal sweep (Fig. 3a). In PD34493, *SF3B1-*K666N was acquired prior to the age of 35 years, whilst *U2AF1*-Q157R initiated clonal growth later (age 41-61) in a previously expanded clade lacking recognisable drivers (Fig. 3b). Interestingly, an additional apparently driverless expansion - a phenomenon recognised in old age^6,32^ - was observed in this individual (Fig. 3b), and a further 3 such expansions in PD41305 (Fig. 3c). In PD41305, the *SRSF2*-P95H mutation was present in only one colony, preventing characterisation of its phylogeny beyond the observation that it was acquired after the age of 13 years (Fig. 3c).

**Fig. 3:**
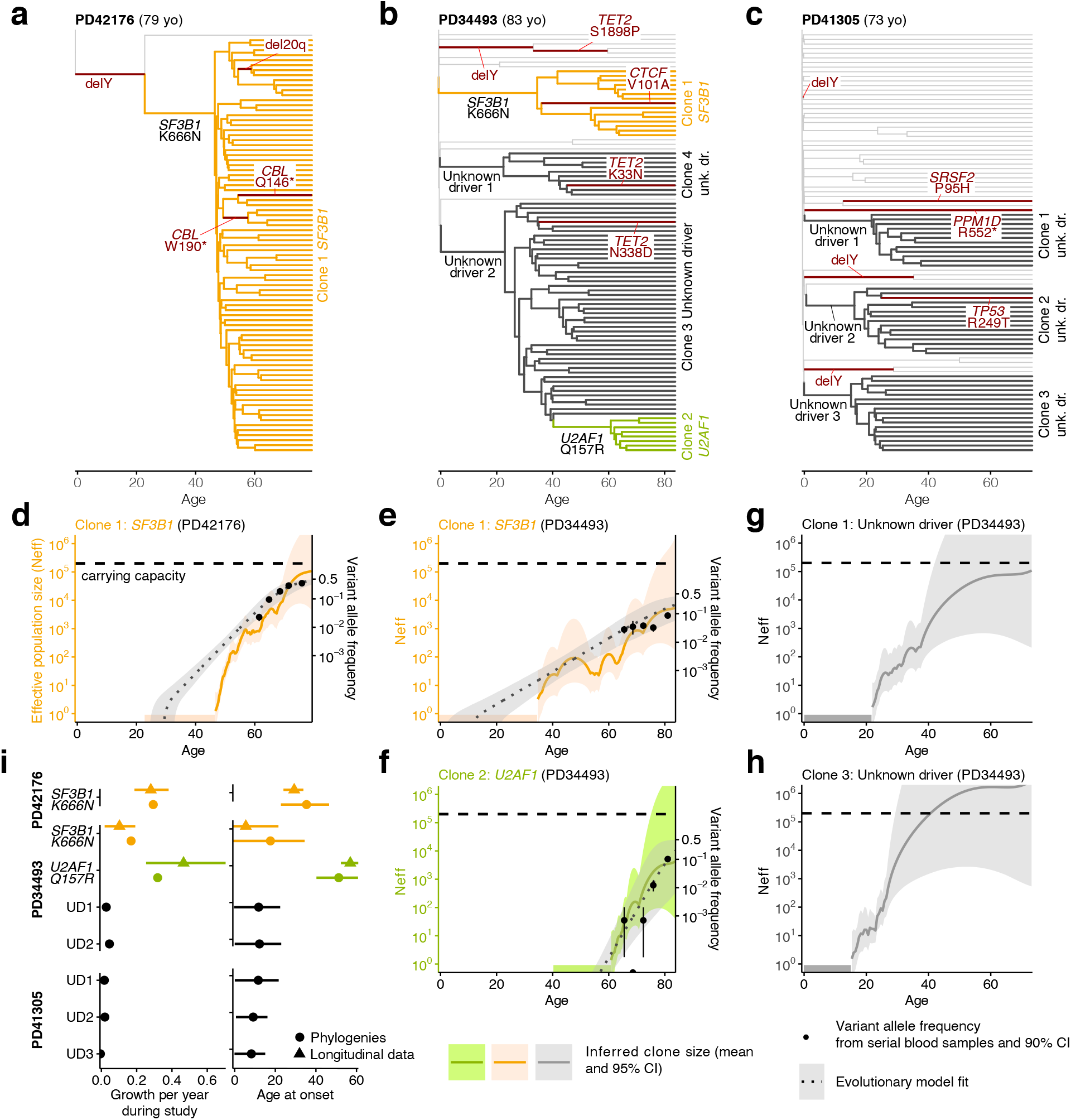
Haematopoietic phylogenetic trees. **a-c**, Haematopoietic phylogenies of participants PD41276 (a), PD34493 (b) and PD41305 (c). Each tree tip is a single cell-derived colony and tips with shared mutations coalesce to an ancestral branch, from which all colonies in such a “clade” arose. Branch lengths are proportional to the number of somatic mutations, which accumulate linearly with age. Branches containing known driver mutations or chromosomal aberrations are annotated. Clonal expansions are coloured: *SF3B1*-K666N-mutant expansions in orange, *U2AF1*-Q157R-mutant expansions in green, and expansions without identified drivers (‘Unknown driver’ or ‘UD’) in black. **d-h**, Growth trajectories of each clonal expansion, as determined by (i) phylogenies (effective population size (Neff) estimated using phylodynamic methods), and (ii) time-series data (using serial VAF measurements and modelled historical growth, as illustrated in Fig. 2, if available). Phylogeny-derived age at clone onset range is represented as a horizontal coloured bar on the x-axis, with the limits of the bar corresponding to the age range of the phylogeny branch along which the corresponding driver mutation was acquired. **i**, Comparison of the ages at onset (right) and growth rate during study period (left) derived from phylogenetic trees and longitudinal data.

We next used the timing and density of clonal branchings (or “coalescences”) to reconstruct the entire growth trajectories of expanded clades using phylodynamic principles (Fig. 3d-h)^29,33,34^. This revealed that the three clades with identified drivers (*SF3B1*-K666N and *U2AF1*-Q157R in PD34493, and *SF3B1*-K666N in PD41276), expanded (Fig. 3d-f) at calculated rates similar to those observed in our time-series VAF measurements during older age (Fig. 3i, left panel). Of note, *SF3B1*-K666N was associated with a substantially different growth rate in PD41276, where it expanded at 28%/yr by serial VAFs (29%/yr by phylodynamic estimate), versus 10%/yr in PD34493 (17%/yr by phylodynamics) (Fig. 3i). Reasons for this difference are unclear, but it is notable that the faster-growing clone had antecedent Y loss (Fig. 3a), an aberration seen in clades from all three individuals and associated with only modest clonal expansion when isolated (Fig. 3a-c). Interestingly, clones without known drivers began to expand within the first two decades of life and grew over their lifetimes at rates comparable to clones with known drivers (14-32%/yr) (Fig. 3g,h, Extended data Fig. 6).

### Many clones decelerate before older age

As the phylodynamic reconstruction of a clone goes back to its inception, we investigated whether clonal growth dynamics during earlier life deviate from the stable growth observed during older age. To corroborate observations from the three individuals depicted in Fig. 3, we conducted additional phylodynamic analyses of trees derived from 1,461 whole-genome sequenced single cell-derived colonies from another four individuals aged 75-81yrs from the study by Mitchell *et al.*^32^. This revealed that, in many instances, the reconstructed effective population size (Neff) of any individual clone grew more slowly towards the sampling date, before it saturated the HSC compartment (Fig. 4a-b; Extended Data Fig. 7a-c). This characteristic deceleration was quantified by fitting a biphasic exponential growth model to early and late parts of the trajectories (Fig. 4c). In most cases, extrapolating early growth (a consistent estimator of the fitness advantage of a clone in Wright-Fisher simulations, Extended Data Fig. 7d, Extended Data Fig. 8) led to dramatic overestimations of clade size (median 35x; Fig. 4d, Extended Data Fig. 7e).

**Fig. 4:**
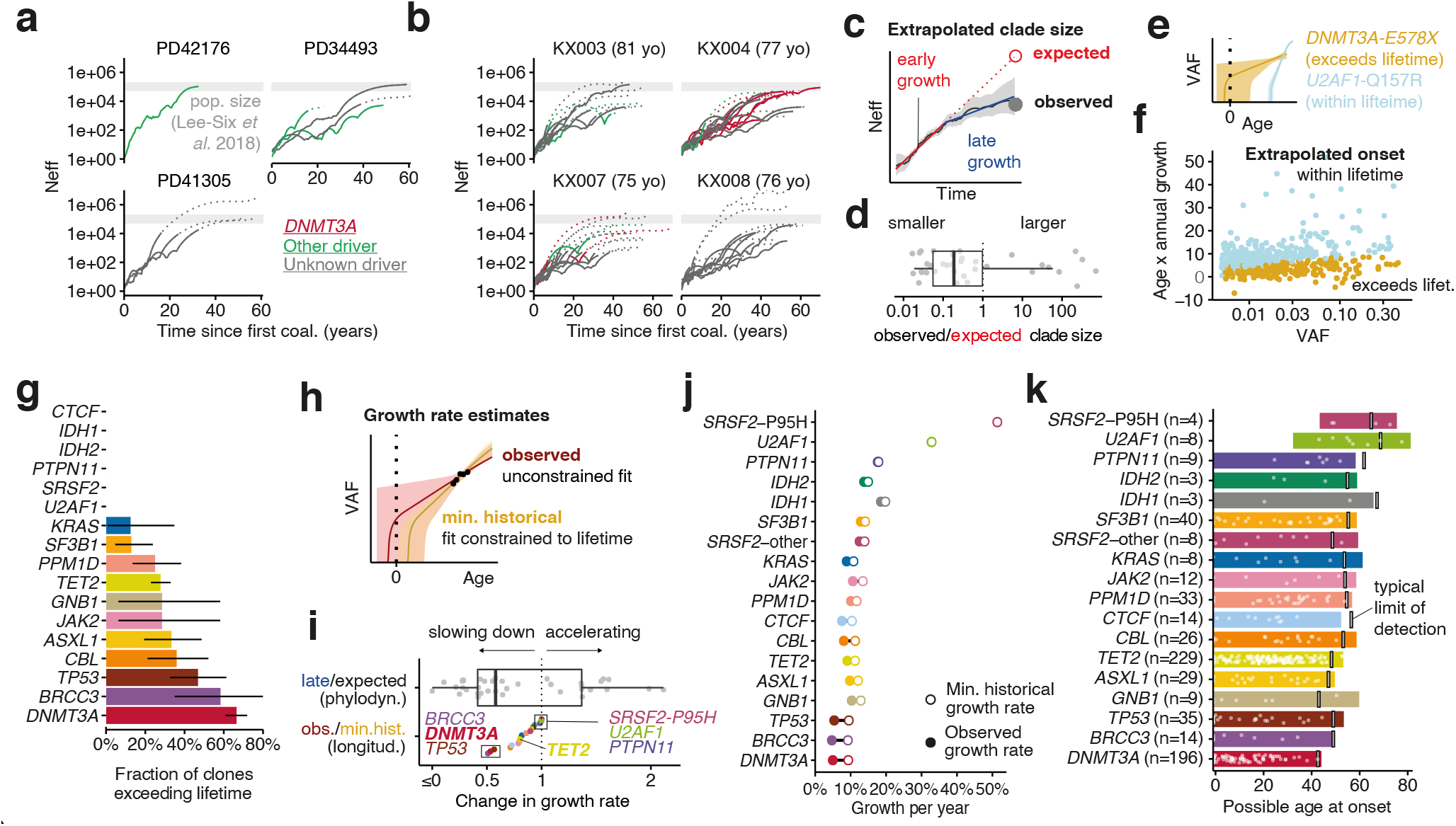
Evidence for clonal deceleration from single-cell phylogenies and longitudinal data. **a,b.**Effective population size (Neff) trajectories inferred from single cell phylogenies in this paper (a) and in Mitchell et al^32^ (b). Dotted lines represent parts of the trajectory with high variance (log(var(Neff)) > 5). **c.** Representation of biphasic fit to Neff estimates and extrapolation from early growth (observed clone size is calculated as the clonal fraction in the phylogeny scaled by an Neff of 200,000 HSC x yr; comparison with 1,000,000 HSC x yr in Extended Data Fig. 7e). **d.** Ratio between observed and expected (extrapolated from early growth) clone size from phylogenies. **e**. Representation of extrapolated trajectories derived from longitudinal data, assuming stable lifelong growth at the same fixed rate we observed during older age; some projections are not feasible (ie. exceeding lifetime, with onset pre-conception). **f**. Relationship between age and the observed growth rate of clones and VAF (longitudinal data; light blue represents clones with projected onset within lifetime and golden represents those exceeding lifetime). **g**. Quantification of unfeasible clones (exceeding lifetime) per gene (longitudinal data). **h.** Representation of the calculation of minimum historical growth. **i.** Quantification of the ratios between observed and historical (longitudinal data) and between late and expected (phylogenetic data) growth. **j.** Differences between the median observed and historical growth per year for each gene. **k.** Projected ages at onset for all clones, assuming stable lifelong growth at the same fixed rate we observed during older age.

We used our longitudinal cohort to orthogonally test the lifelong stability of clonal growth by extrapolating the observed (fitted) trajectory of each clone backwards in time to infer the age at clonal onset. To account for stochastic drift, which can lead to faster growth of small clones, and the finite carrying capacity of the HSC population, which naturally limits/slows large clones, we derived and used an approximation to a Wright-Fisher process (Extended Data Fig. 4a,b). While estimates of age at clonal onset agreed with phylogenetic estimates for the fast-growing splice factor mutations (Fig. 3i), for many other clones, constant lifelong growth at the rate we observed during old age would be too slow to explain the observed VAFs (Fig. 4e,f,g), proposing that clonal expansion was faster in earlier life. These observations reveal that, at least for some clones/genes, the dynamics observed in later life are not representative of those that prevail earlier.

We then assessed the minimum lifetime rate at which clones must have grown in order to reach the observed VAFs in our longitudinal data, henceforth termed ‘historical growth’, by restricting fits/solutions to growth rates that would place the age of clonal onset within individuals’ lifetimes (Fig. 4h, Supplementary Table 8). Expectedly, this minimal historical growth rate was typically higher than the growth rate observed during the study period (i.e. in older age; Fig. 4i, Extended Data Fig. 7f). Moreover, the fold-changes between historical and observed growth rates derived from longitudinal data were qualitatively in good agreement with the fold-changes between late growth and expected growth (the latter assuming growth is constant through life and carrying capacity is fixed) derived from phylodynamic data (Fig. 4c,i, Extended Data Fig. 7f). Taken together it thus emerges that many clones grew more rapidly early in life compared with the rate in old age.

### Driver-specific differences in lifetime clonal behaviour

The effect of deceleration was most marked for clones bearing mutations in *DNMT3A*, *BRCC3* and *TP53*, whose early growth was at least twice as fast as that measured during old age (Fig. 4i,j). Conversely, we observed almost no deceleration of fast-growing clones harbouring *U2AF1*, *SRSF2*-P95H, *PTPN11* or *IDH1* mutations (Fig. 4i,j). It is particularly notable that the *TET2*-mutant clones were much less susceptible to deceleration than *DNMT3A-*mutant clones (Fig. 4i-j). This is consistent with the observation that the prevalence of *TET2*-mutant CH rises at older ages and eventually exceeds that of *DNMT3A*-mutant CH, which is more prevalent at younger ages (Fig. 1d). A declining relative advantage of *DNMT3A* mutations in older age was also suggested by the much lower proportion of *DNMT3A* mutant-clones reaching detectable limits during our study period compared to clones bearing mutations in other genes (“incipient clones”, Extended Data Fig. 9a).

To derive representative ranges for age at clone onset for each driver gene, we capped individual estimates at conception, thus avoiding estimates that projected beyond individuals’ lifetimes (Fig. 4k, Extended Data Fig. 9b,c). We also validate this method using simulations and confirm that these ranges are not affected by changes in Neff or generation time (Extended Data Fig. 9d,e). We estimated that the average latency between clone foundation and detection in peripheral blood at VAF ≥0.2% (Supplementary Note 1) was 30 years across all clones, with considerable variability between mutant genes, ranging from 38 years for *DNMT3A*-mutant clones to 12 years for *U2AF1*-mutant clones. Most drivers were projected to initiate expansions of clones throughout life, compatible with the notion that somatic mutations occur at a constant rate^28,29,35^. However, solutions for *DNMT3A*-mutant clones concentrated earlier in life, consistent with early initiation and rapid expansion followed by marked deceleration then slow growth, as discussed earlier. Of note, capping onset at conception is arbitrary and it remains possible that some clones start later and exhibit faster initial growth followed by even stronger deceleration, a scenario that would be more consistent with published fitness estimates of 11-19%/yr based on cross-sectional VAF measurements^12^. In contrast, *SRSF2-*P95H and *U2AF1* mutations initiated clonal expansion always after 30 years of age and with a median age at onset of 58 and 57 years, respectively (Fig. 4k). This indicates that the reported rarity of these mutant clones in people aged <60 years^5,6,8^ is not due to slow growth over decades, but rather due to their late onset followed by rapid expansion and also provides a plausible explanation for the high risk of leukaemic progression associated with these mutations^9,36^.

### CH dynamics and malignant progression

To investigate the links between mutation fitness and malignant progression, we built on our previous study of AML risk prediction^9^ and revealed that among CH driver genes a faster growth rate was associated with a higher AML risk (adjusted R^2^=0.55, p=0.0037, Fig. 5a). For example, genes driving fast CH growth like *SRSF2* and *U2AF1* were associated with the highest risks of leukaemogenesis, while slow-growing clones such as those bearing *DNMT3A* mutations, conferred a lower risk. To confirm our findings in larger studies and include myeloid malignancies other than AML, we analysed large published datasets of AML (n=1540)^37^ and myelodysplastic syndromes (MDS, n=738)^38^ using a site-specific extension of the dNdScv algorithm to formally quantify the extent to which individual hotspots are under the influence of positive selection in these cancers (Supplementary Tables 9,10)^25^. This analysis revealed a positive correlation between each hotspot’s growth coefficient in CH and its selection strength in myeloid cancer (Fig. 5b; adjusted R^2^=0.19, p=0.0016), corroborating the AML risk analysis. Nevertheless, the observation that the same CH driver gene can progress to either AML or MDS, with variable predilections as quantified by gene-level dN/dS comparison (Extended Data Fig. 10; Supplementary Table 10), suggests that factors other than growth rate can also influence a mutation’s malignant potential.

**Fig. 5:**
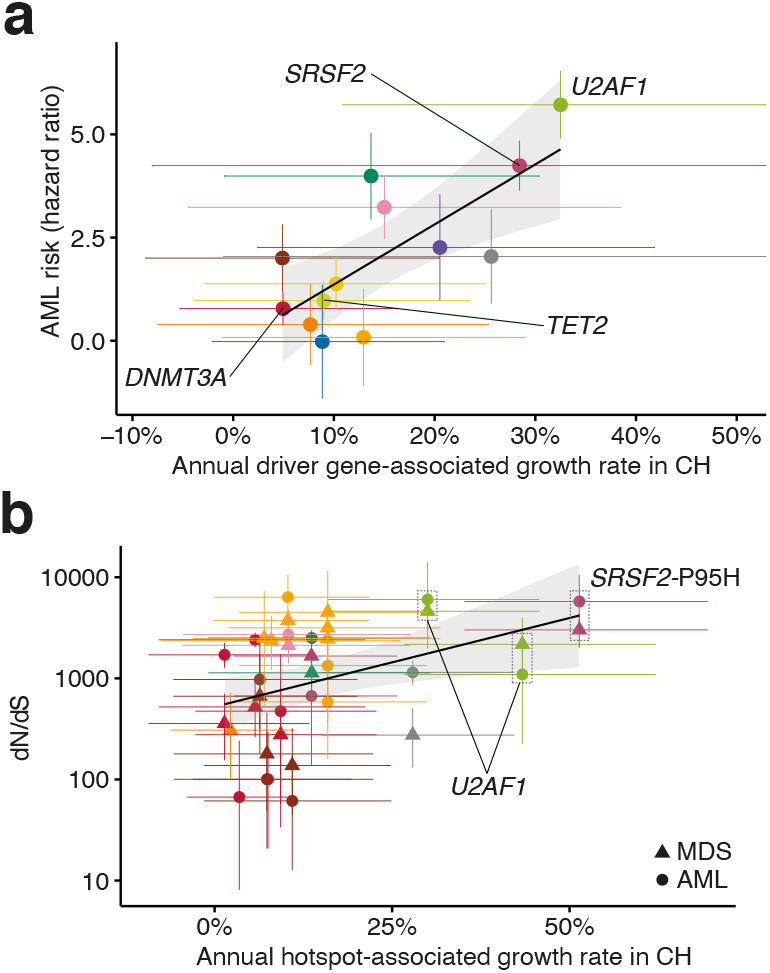
CH dynamics and progression to myeloid disease. **a**, Relationship between the growth rate associated with each driver gene in CH, and the risk of AML progression associated with that driver gene. **b**, Relationship between the growth rate associated with each recurrent mutation in CH, and the strength of selection of that mutation in AML (circles) and MDS (triangles). In **a** and **b** genes/hotspots mentioned in the main text are highlighted.

## Discussion

The phenomenon of CH has served as an exemplar in the developing understanding of somatic mutation, clonal selection and oncogenesis in human tissues^4,39^. However, the nature of these interrelated processes can change over time and their consequences develop only slowly, making them difficult to investigate. Here, we studied the longitudinal behaviour of CH over long periods (median 13 years) and combined this with lifelong phylodynamic analyses of haematopoiesis to derive new insights into these fundamental biological processes.

First, we found that most clones (92%) display stable exponential growth dynamics in older age, at rates influenced by their driver mutations. This allowed us to predict future clonal growth trajectories, a finding with potentially useful implications for clinical practice (Extended Data Fig. 4e-g). Surprisingly, mutations in *DNMT3A*, reportedly the most common CH driver gene^5–7^, were associated with slower clonal expansion than most other CH genes. Also, *DNMT3A* hotspot mutations (e.g. at codon R882) were not associated with faster growth than other *DNMT3A* mutations (Figure 2c). By contrast, *TET2-*mutant clones expanded significantly faster over the study period (Fig. 2c) and, reflecting this, also reached detectable levels much more frequently on-study than *DNMT3A*-mutant clones (Extended Data Fig. 9a). This resulted in *TET2* becoming the most prevalent CH driver after the age of 75 years (Figure 1d).

These initial findings suggested that, while clonal growth is remarkably stable in old age, dynamics in earlier life may deviate from this behaviour, challenging the premise that mutation fitness is constant over the human lifespan^12^. To test this, we first attempted to derive when individual CH clones were founded, using simple retrograde extrapolation of observed trajectories. This led to projected ages at clonal foundation that preceded conception for a large number of clones (Fig. 4f,g), implying that their early growth must have been faster than that we observed during old age. This was most striking for *DNMT3A*, for which more than two thirds of projections were implausible (ie. onset pre-conception), but less common for *TET2* and very uncommon for splicing factor genes (Fig. 4g).

To further investigate lifelong clonal behaviour, we analysed haematopoietic phylogenies from healthy old individuals and found that aged haematopoiesis was dominated by a small number of expanded HSC clones, some of which lacked recognisable drivers^32^. Using phylodynamic approaches to track clonal growth rates through life, in conjunction with findings from our longitudinal cohort, we reveal widespread clonal deceleration prior to the period of stable growth during old age, in the context of an increasingly competitive oligoclonal HSC compartment (Fig. 4i). *DNMT3A*-mutant clones, as well as those bearing mutations in *TP53* and *BRCC3* and also apparently driverless clones, were among those displaying the most marked degree of deceleration (Fig. 4i). In contrast, *TET2* mutations appeared to drive more stable lifelong growth (Fig. 4h-j), which may underlie their apparent ability to initiate clonal expansion fairly uniformly through life (Fig. 4k) and the fact that *TET2* “overtakes” *DNMT3A* as the most common CH driver after the age of 75 years (Fig. 1d).

In diametric contrast to *DNMT3A* and unlike other genes, CH driven by mutant *U2AF1* and *SRSF2*-P95H only initiated late in life (Fig. 4k) and exhibited some of the fastest expansion dynamics (Fig. 2c). These data were corroborated by phylogenetic analyses (Fig. 3b,f) and tally with the sharp increase in prevalence of splice factor-mutant CH^8^, MDS^38,40,41^ and AML^37,42^ in old age and the high risk of progression to myeloid cancers associated with these mutations^9^. The particular behaviour of these clones proposes a specific interaction with ageing, which could relate to cell-intrinsic factors or to cell-extrinsic changes in the aging haematopoietic niche that make it more suitable for HSCs harbouring splice factor mutations^43,44^.

Finally, we explored the relationship between clonal growth rate in CH and the development of myeloid cancers. We find that mutations associated with faster CH growth are also those associated with higher risk of progression to AML (Fig. 5a) and are under the strongest selective pressure in AML and MDS (Fig. 5b). Indeed, we show that the average annual growth per gene explains over 50% of the variance in AML risk progression. This shows that an improved understanding of growth dynamics in CH can help identify those at risk of myeloid malignancies.

Collectively, our work gives new insights into the lifelong clonal dynamics of different subtypes of CH, the impact of ageing on haematopoiesis, and the processes linking somatic mutation, clonal expansion and malignant progression.

## Methods

### Study participants

Ethical permission for this study was granted by The East of England (Essex) Research Ethics Committee (REC reference 15/EE/0327). The SardiNIA longitudinal study recruited individuals from four towns in the Lanusei Valley in Sardinia, capturing 5 phases of sample and data collection over more than 20 years^25^. We analysed serial samples from 385 individuals in the SardiNIA project.

### Targeted sequencing and variant-calling

Target enrichment of genomic DNA was performed using a custom RNA bait set (Agilent SureSelect ELID 3156971), designed complementary to 56 genes implicated in CH and haematological malignancies (Supplementary Table 1). Libraries were sequenced on Illumina HiSeq 2000 and variant-calling was performed as we described previously^9,45^. Briefly, somatic single-nucleotide variants and small indels were called using Shearwater (v.1.21.5), an algorithm designed to detect subclonal mutations in deep sequencing experiments^46^. Two additional variant-calling algorithms were applied to complement this approach: CaVEMan (v.1.11.2) for single-nucleotide variants, and Pindel (v.2.2) for small indels^47,48^. VAF correction was performed using an in-house script (https://github.com/cancerit/vafCorrect). Finally, allele counts at recurrent mutation hotspots were verified using an in-house script (github.com/cancerit/allelecount). Variants were filtered as we described previously^9,45^, but were not curated with regard to existing notions of oncogenicity, ie. all somatic variants passing quality filters were retained for analysis.

If a variant was identified in an individual at any time-point in the study, this site was re-queried in the same individual at all other time-points, using an in-house script (cgpVAF) to provide pileup (SNV) and Exonerate (indel) output (https://github.com/cancerit/vafCorrect). No additional filters were applied to these back-called variants.

### Selection analyses (dN/dS)

To quantify selection, we used the dNdScv algorithm, a maximum-likelihood implementation of dN/dS, which measures the ratio of non-synonymous (N) to synonymous (S) mutations, while controlling for gene sequence composition and variable substitution rates^26^. We first applied this method to the mutation calls from the longitudinal SardiNIA cohort in order to identify which genes are under positive selection in the context of CH. For this analysis, any mutation that was present in a single individual at multiple time-points was counted only once.

To characterise patterns of selection in AML and MDS, we applied dNdScv to two published data sets. The AML set was derived from 1540 patients enrolled in three prospective trials of intensive therapy^37^. The MDS set included 738 patients with MDS or closely related neoplasms such as chronic myelomonocytic leukaemia^38^. Both used deep targeted sequencing of 111 cancer genes, which overlapped with 13 of the 17 genes of interest in our longitudinal CH study (*PPM1D, CTCF, GNB1* and *BRCC3* were not sequenced in the AML/MDS studies). We called and filtered variants in the 13 overlapping genes using the strategy described above (under ‘Targeted sequencing and variant-calling’). Variants were identified in all 13 genes in both AML and MDS datasets (Supplementary Table 10). We calculated dN/dS values both at the level of individual genes, and at single-site level for hotspots, the latter using the sitednds function in the dNdScv R package.

### Hierarchical modelling of clone trajectories through time

We use Bayesian hierarchical modelling to model clonal trajectories. Since we are unable to phase different mutations into specific clones and given that individual CH clones typically harbour a single mutation^49^, we assume that each mutation is heterozygous and its VAF is representative of the prevalence of a single clone. Accordingly, for a given individual *j* and mutation *i*, we have a mutant clone *c*_*ij*_. We model the counts 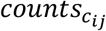 for *c*_*ij*_ at age *t* as a binomial distribution, such that 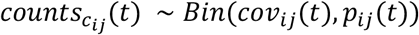, with *cov*_*ij*_ as the coverage of this mutation at age *t* and *p*_*ij*_(*t*) ~ *Beta*(*α*(*t*), *β*) as the expected proportion of mutant allele copies. As such, 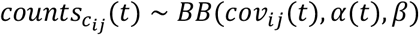. Here, *β* ~ *N*(*μ*_*od*_, *σ*_*od*_) is the technical overdispersion whose parameters are estimated using replicate data (details below) and 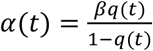, where 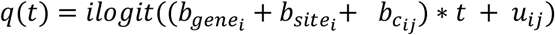. We use this parameterization to guarantee that 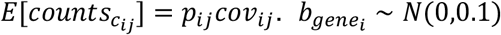 and 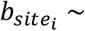 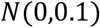 are the gene and site growth effects for mutation *i*, respectively. 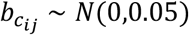 is the growth effect associated exclusively with mutation *i* in individual *j* - i.e. of mutant clone *c*_*ij*_ - and *u*_*ij*_ is the offset accounting for the onset of different clones at different points in time. We also define the growth effect of *c*_*ij*_as 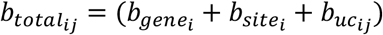. Along this work we will refer to 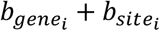 as the *driver (growth) effect* and to 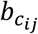 as the *unknown-cause (growth) effect* - the fraction of growth that is quantifiable but not explained by either gene or site.

#### Preventing identifiability issues and reducing uninformed estimates

To address possible identifiability issues in our model, when a *gene* has a single mutation (*JAK2-*V617F and *IDH2*-R140Q), the effect is considered to occur only at the *site* level. To avoid estimating the dynamics of a site from a single individual, we only model 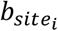 when two or more individuals have a missense mutation on site *i* - we refer to these sites as “recurrent sites”. Overall, we consider a total of 17 genes and 39 recurrent sites (Supplementary Table 5).

#### Estimating and validating growth parameters

Using the model described above, we use Markov Chain Monte Carlo (MCMC) with a Hamiltonian Monte Carlo (HMC) sampler with 150-300 leapfrog steps as implemented in greta^50^. We sample for 5,000 iterations and discard the initial 2,500 to get estimates for the distribution of our parameters. As such, our estimates for each parameter are obtained considering their mean, median and 95% highest density posterior interval for 2,500 samples.

We assess the goodness-of-fit using the number of outliers detected in any trajectory and consider only trajectories with no outliers as being explained by our model and, as such, growing at constant rate. Outliers are assessed by calculating the tail probabilities of the counts under our model with a hard cut-off at 2.5%. As such, *P*_*outlier*_ = 1 if 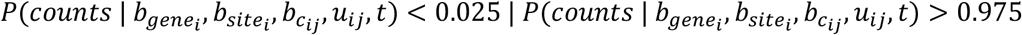 and *P*_*outlier*_ = *0* otherwise. We validate this approach using Wright-Fisher simulations (Supplementary Methods). We additionally assess the predictive power of this model on an additional time-point that was available for a subset of individuals and that was not used in the inference of parameters in our model (Supplementary Methods).

#### Estimating the technical overdispersion parameter

Technical VAF overdispersion used two distinct sets of data:

1. Horizon Tru-Q-1 was serially diluted to VAFs of 0.05, 0.02, 0.01, 0.005 and 0 using Horizon Tru-Q-0 (verified wild-type at these variant sites), then sequenced in duplicate or triplicate;
2. 19 SardiNIA samples with mutations across 15 genes at a range of VAFs, were sequenced in triplicate.

Sample processing and analysis was performed as described in the “Targeted Sequencing and Variant-calling” section. Replicate samples were picked from the same stock of DNA, then library preparation and sequencing steps were performed in parallel. Variant calls for these replicate samples are in Supplementary Table 11.

For (1), we model the distribution over the expected *VAF* as a beta distribution such that *VAF* ~ *Beta*(*α*, *β*) and for (2) we adopt a model identical to the one described earlier in this section but use only gene growth effects 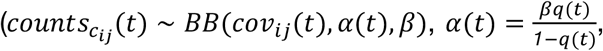 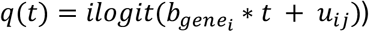. Here, we model *β* ~ *exp*(*r*) with *r* as a variable with no prior. We use MCMC with HMC sampling with 400-500 leapfrog-steps as implemented in greta^50^ to estimate the mean and standard deviation of *β*. For this estimate we use 1,000 samples from the posterior distribution.

### Analysis of non-mutation factors as determinants of clonal growth rate

#### Inherited polymorphisms and JAK2-mutant clonal growth

The SardiNIA cohort had previously been characterised using two Illumina custom arrays: the Cardio-MetaboChip and the ImmunoChip^25^. Inherited genotypes at 12 loci previously associated with MPN risk were extracted for the 12 individuals with *JAK2*-V617F mutation^22,23^. The relationship between each individual’s total number of inherited risk alleles and *JAK2*-mutant clonal growth rate was assessed by Pearson’s correlation. The 46/1 haplotype, which harbours 4 SNPs in complete linkage disequilibrium, was considered as a single risk allele.

#### Age, sex and smoking experience

We assess the association between unknown-cause growth and age through the calculation of a Pearson correlation considering all genes, both together and separately while controlling for multiple testing. We also assess the association between unknown-cause growth and sex and smoking history using a multivariate regression where unknown-cause growth is the dependent variable and sex and previous smoking experience are the covariates, while also controlling for age.

### Determining the expected age at beginning of clone onset

We consider that HSC clones grow according to a Wright-Fisher model. According to this, for an initial population of HSC *n*/*2*, we can consider two scenarios - that of a single growth process where the time at which the cell first starts growing *t*_*0*_ is described as 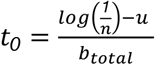 or that of a two step growth process, where 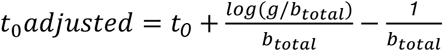, where *g* is the number of generations per year. The latter scenario is the one chosen, due to its strong theoretical foundation and previous application to mathematical modelling of cancer evolution^51^. The two regimes that describe it are an initial stochastic growth regime and, once the clone reaches a sufficient population size, a deterministic growth regime. The adjustment made to *t*_*0*_ in *t*_*0*_*adjusted* can be interpreted as first estimating the age at which the clone reached the deterministic growth phase 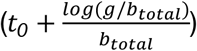 followed by subtracting the expected time for a clone to overcome its stochastic growth phase 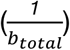. For both *n* and *g* we use the estimates based on ^29^ - *n* = *50,000* and *g* = *2*. We validate this approach using simulations (Supplementary Methods) and test the approach against our serial VAF data and verify that changes in *n* and *g* do not have a dramatic impact on age at onset estimates by considering a range of values (*n* = {*10,000*; *50,000*; *100,000*; *200,000*; *600,000*} and *g* = {*1*; *2*; *5*; *10*; *13*; *20*}).

### Derivation of blood colonies and phylogenetic tree construction

#### Sample preparation and sequencing

We selected 3 individuals with splicing gene mutations from the SardiNIA cohort for detailed blood phylogenetic analysis. Peripheral blood samples were drawn into Lithium-heparin tubes (vacutest, kima, 9ml) and buccal samples were taken (Orangene DNA OG-250). Peripheral blood mononuclear cells were isolated from blood and plated at 50,000 cells per ml in MethoCult 4034 (Stemcell Technologies). After 14 days in culture, 96 single haematopoietic colonies were plucked per individual (total 288 colonies) and lysed in 50μl of RLT lysis buffer (Qiagen).

Library preparation for whole genome sequencing (WGS) was performed using our low-input pipeline as previously described^52,53^. 150bp paired-end sequencing reads were generated using the NovaSeq® 6000 platform to a mean sequencing depth of 15x per sample. Reads were aligned to the human reference genome (NCBI build37) using BWA-MEM.

#### Variant-calling and filtering

Single-nucleotide variants (SNVs) and small indels were called against an unmatched reference genome using the in-house pipelines CaVEMan and Pindel, respectively^47,48^. ‘Normal contamination of tumour’ was set to 0.05; otherwise standard settings and filters were applied. For all mutations passing quality filters in at least one sample, in-house software (cgpVAF, https://github.com/cancerit/vafCorrect) was used to produce matrices of variant and normal reads at each mutant site for all colonies from that individual. Copy-number aberrations and structural variants were identified using matched-normal ASCAT^54^ and BRASS (https://github.com/cancerit/BRASS). Low-coverage samples (mean <4x) were excluded from downstream analysis (n=1, PD41305). Samples in which the peak density of somatic mutation VAFs was lower than expected for heterozygous changes (in practice VAF<0.4) were suspected to be contaminated or mixed colonies, and were also excluded from further analysis (n=3, PD41305; n=9, PD41276; n=3, PD34493).

Multiple post-hoc filtering steps were then applied to remove germline mutations, recurrent library prep / sequencing artefacts, and in vitro mutations, as described previously^55^ and detailed in custom R scripts (https://github.com/margaretefabre/Clonal_dynamics). Buccal samples were used as an additional filter; mutations were removed if the variant:normal count in the buccal sample was consistent with that expected for a germline mutation (0.5 for autosomes and 0.95 for XY chromosomes, binomial probability >0.01), and were retained if (i) the variant:normal count in the buccal sample was *not* consistent with germline (binomial probability <1×10^−4^) and (ii) the mutation was not present in either of 2 large SNP databases (1000 Genomes Project and Kaviar) with MAF > 0.001.

#### Phylogenetic tree construction and assignment of mutations back to the tree

These steps were also performed as described previously^55^ and are detailed here:

https://github.com/margaretefabre/Clonal_dynamics. Briefly, sample were assigned a genotype for each mutation site passing filtering steps (‘present’ = ≥2 variant reads and probability > 0.05 that counts came from a somatic distribution; ‘absent’ = 0 variant reads and depth ≥6; ‘unknown’ = neither ‘absent’ criteria met). A genotype matrix of shared mutations was fed into the MPBoot program^56^, which constructs a maximum parsimony phylogenetic tree with bootstrap approximation. The in-house-developed R package treemut (https://github.com/NickWilliamsSanger/treemut), which uses original count data and a maximum likelihood approach, was then utilised to assign mutations back to individual branches on the tree. Since individual edge length is influenced by the sensitivity of variant-calling, lengths were scaled by 1/sensitivity, where sensitivity was estimated using the proportion of germline variants called.

#### Reconstruction of population trajectories

Phylogenies were made ultrametric (branch lengths normalised) using a bespoke R function (make.tree.ultrametric, https://github.com/margaretefabre/Clonal_dynamics/my_functions). Assuming a constant rate of mutation acquisition^28,29,35^, the time axis was scaled linearly, where the root of the tree represents conception, and the tips represent age at sampling. We then analysed population size trajectories by fitting Bayesian nonparametric phylodynamic reconstructions (BNPR) as implemented in the phylodyn R package^33,34^ to clades - sets of samples in a phylogenetic tree sharing a most recent common ancestor (MRCA) - defined by either having a driver mutation on the MRCA or a MRCA branch length that spans more than 10% of the tree depth and with 5 tips or more. We also estimated the lower and upper bounds for age at onset of clonal expansion to be the limits of the branch containing the most recent common ancestor.

### Deceleration in phylogenies and longitudinal data

We detect deceleration using two different approaches - the ratio between expected and observed clone size using phylodynamic estimates and the ratios between observed and historical (from longitudinal data) and between late and expected (from phylogenetic data), respectively. To obtain the late growth rate we fit a biphasic log-linear model to our phylodynamic estimation of Neff - this enables us to obtain an *early* and a *late* growth rate (details in the Supplementary Methods).

#### Expected and observed clone size

The expected clone size is calculated by extrapolating the early growth rate until the age of sampling; having this we can calculate the ratio between expected and observed growth. The ratio between these quantities is then used as a measure of deceleration (details in the Supplementary Methods).

#### Growth ratio in phylogenetic data

The late growth rate is defined as the late growth rate defined in the previous section of the methods. The expected growth rate for the phylogenies is calculated as the growth coefficient for a sigmoidal regression that assumes a population size of 200,000 HSC as the carrying capacity. We then use the ratio between these quantities as a measure of deceleration (1 implies no deceleration; <1 implies deceleration).

#### Growth ratio in longitudinal data

The observed growth rate is defined as the growth rate inferred directly from the data. The minimal historical growth is the growth rate estimate obtained by restricting clone initiation to a time after conception (age at at onset > −1).

### Associations between CH dynamics and (i) AML progression and (ii) selection in MDS and AML

To calculate the association between CH dynamics and AML we used the risk coefficients from our previous work in predicting the onset of AML^9^, which were calculated by fitting a Cox-proportional hazards model that calculated the risk of AML onset associated with each gene while controlling for age, sex and cohort, and estimate the coefficient of correlation between the expected value of the annual growth for the posterior distribution of each gene (considering gene, site and unknown-cause effects) and the AML progression risk.

The association between CH dynamics and selection in MDS and AML use the dN/dS values calculated with dNdScv as previously described in the methods, using two distinct cohorts from previous studies^37,38^. dN/dS values were calculated for all hotspots and their coefficient of correlation with the expected value of the annual growth for the posterior distribution of each hotspot (also considering gene, site and unknown-cause effects) was calculated.

### Statistical analyses

All statistical analyses were conducted using the R software^57^ - MCMC models were fitted using gret^50^ and hypothesis testing, generalised linear models and maximum likelihood fits were performed in base R.

## Acknowledgements

This work was funded by a joint grant from the Leukemia and Lymphoma Society (RTF6006-19) and the Rising Tide Foundation for Clinical Cancer Research (CCR-18-500) and by the Wellcome Trust (WT098051). M.F. is funded by a Wellcome Clinical Research Fellowship (WT098051). J.G.A. is supported by the NIHR Cambridge BRC and their opinions are not necessarily those of the NHS, the NIHR or the Department of Health and Social Care. G.S.V. is funded by a Cancer Research UK Senior Cancer Fellowship (C22324/A23015) and work in his lab is also funded by the European Research Council, Kay Kendall Leukaemia Fund, Blood Cancer UK and the Wellcome Trust. E.F.M. is supported by the Wellcome Trust and Beit Foundation (104064/Z/14/Z) and by the EC H2020. The collection of samples and data from the SardiNIA longitudinal cohort study was supported by the Intramural Research Program of the NIH, National Institute on Aging (NIA) of the National Institute of Health (NIH) with contracts N01-AG-1-2109 and HHSN271201100005C; and by the European Union’s Horizon 2020 Research and Innovation Programme under grant agreement 633964 (ImmunoAgeing).

## Data availability

The data files necessary to run the analysis in https://github.com/josegcpa/clonal_dynamics are freely available at https://doi.org/10.6084/m9.figshare.15029118. All sequencing data have been deposited in the European Genome-phenome Archive (EGA) (https://www.ebi.ac.uk/ega/). Targeted sequencing data have been deposited with EGA accession numbers EGAD00001007682 and EGAD00001007683; WGS data have been deposited with accession number EGAD00001007684. Data from the EGA are accessible for research use only to all bona fide researchers, as assessed by the Data Access Committee (https://www.ebi.ac.uk/ega/about/access). Data can be accessed by registering for an EGA account and contacting the Data Access Committee.

## Code availability

All analyses reported in this study used the statistical software R (v.3.6.3). All R files used for the longitudinal and phylodynamic modelling and validation are publicly available at https://github.com/josegcpa/clonal_dynamics. All files used for the construction of phylogenetic trees are publicly available at https://github.com/margaretefabre/Clonal_dynamics.

## Author contributions

GV and MG conceived and supervised the study. MF, JGA, MSV carried out analyses and generated data figures. MG and JGA developed and implemented the statistical modelling of clonal dynamics. VO, EF, MM and FC oversaw the SardiNIA cohort. VO, EF, MM, EMcK and FC provided samples and data from the Immunoageing study. AD, JR, CH, JB, MF and GV processed participant samples and performed assays. FA, NW, JN and IM generated computational code used in this paper. EM, MC and PC provided single-cell-derived colony WGS data and helped with data analysis/interpretation.

## Extended data figures

**Extended Data Fig. 1:**
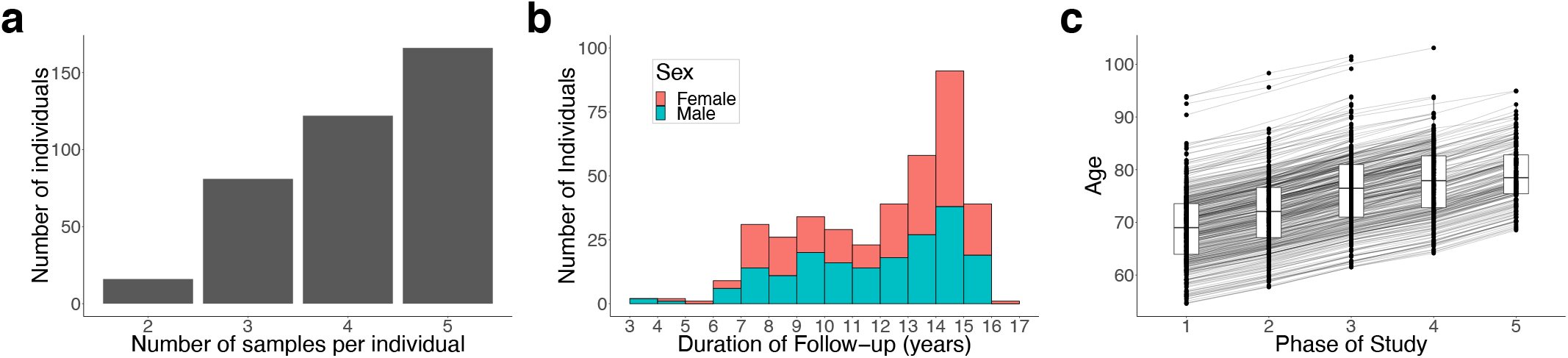
Longitudinal cohort characteristics. **a**, Distribution of the number of serial samples obtained per individual. **b**, Duration of follow-up per individual. **c**, Distribution of participants’ ages at each of the five sampling phases of the SardiNIA study.

**Extended Data Fig. 2:**
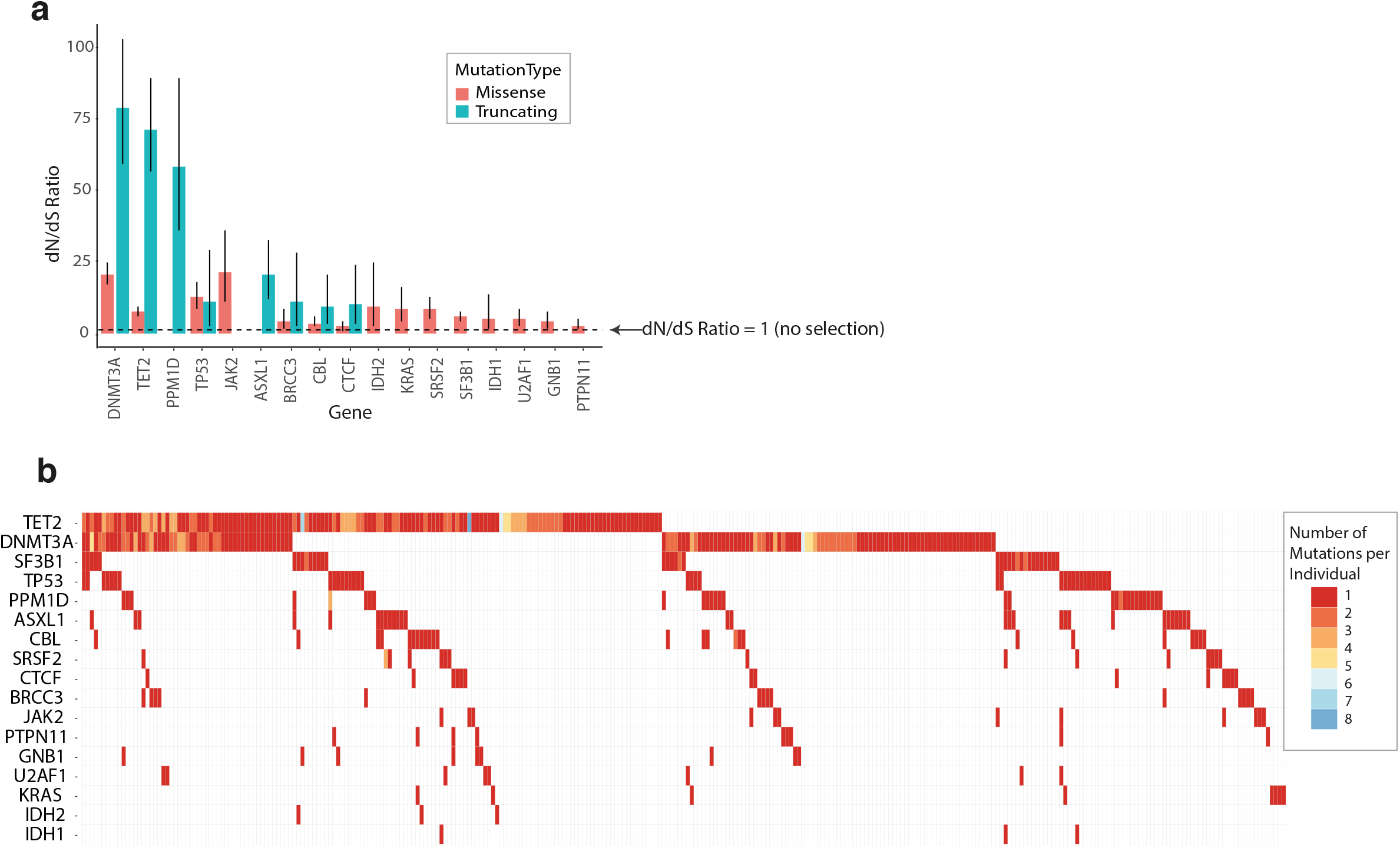
Mutation prevalence and selection in different genes. **a**, Observed-to-expected (dN/dS) ratios for the 17 genes with missense and/or truncating mutations under positive selection (with q<0.1). The dashed line indicates a dN/dS value of 1, which represents neutrality (no selection). **b**, Waterfall plot showing the number and distribution of mutations among participants. Each column represents 1 individual, and each row 1 gene. Coloured squares indicate the presence of a mutation with the specific colour indicating the number of distinct mutations in that gene identified in that individual. For individuals with the same mutation identified at multiple serial time-points, the serially-observed mutation is counted only once.

**Extended Data Fig. 3:**
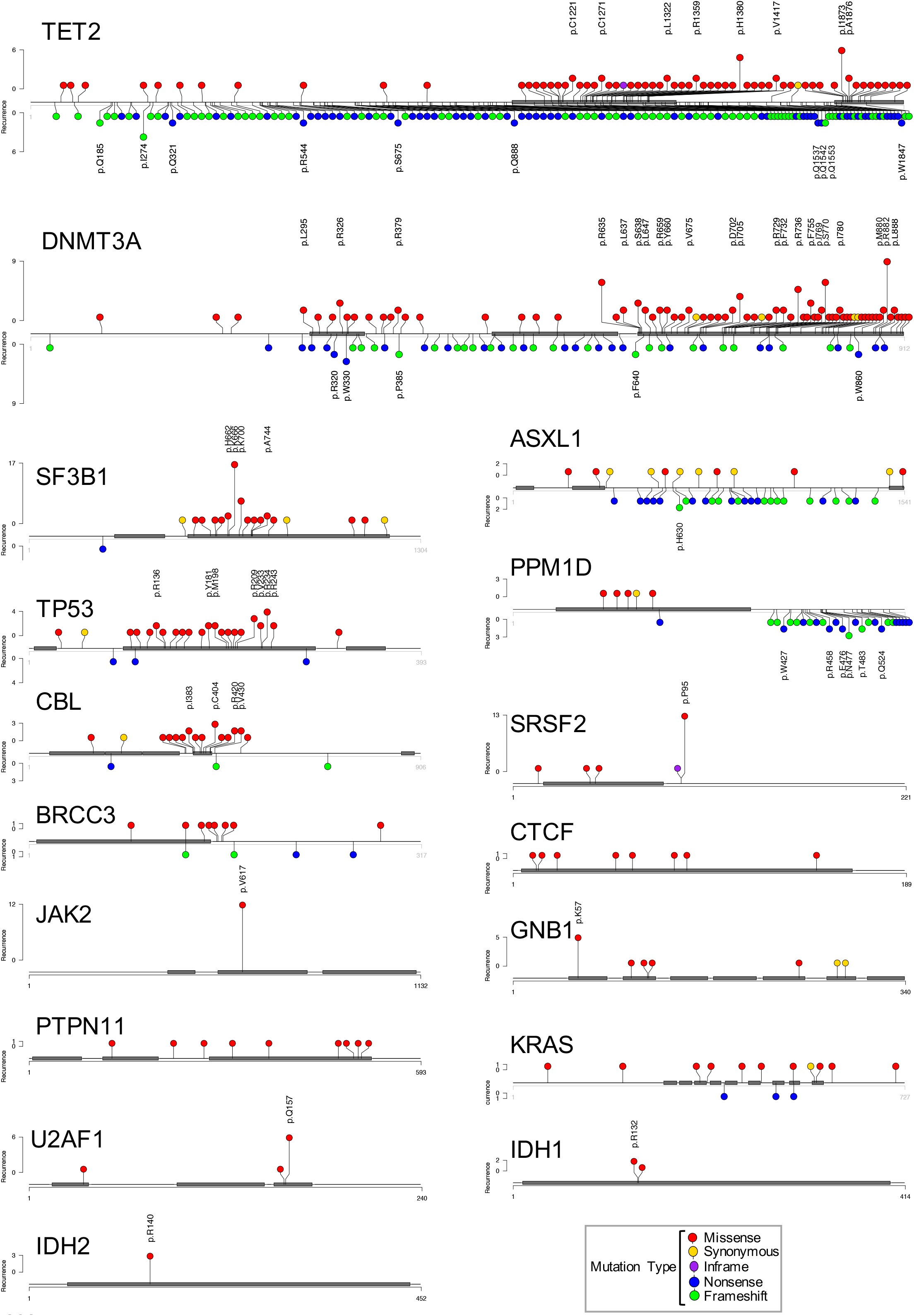
Distribution of somatic mutations within driver genes (previous page). Lolliplots show the longest protein isoform of each gene, with protein domains depicted by grey rectangles. Each circle represents a somatic mutation. The vertical distance of the circle from the protein cartoon indicates its recurrence in the cohort (quantified on the y-axis). Amino acid codons recurrently mutated (ie. observed in more than one individual) in our cohort are explicitly labelled. Circle colours indicate the mutation type as per key. Non-truncating mutations (missense, inframe, synonymous) are depicted above and truncating mutations (nonsense, frameshift) below the protein cartoon.

**Extended Data Fig. 4:**
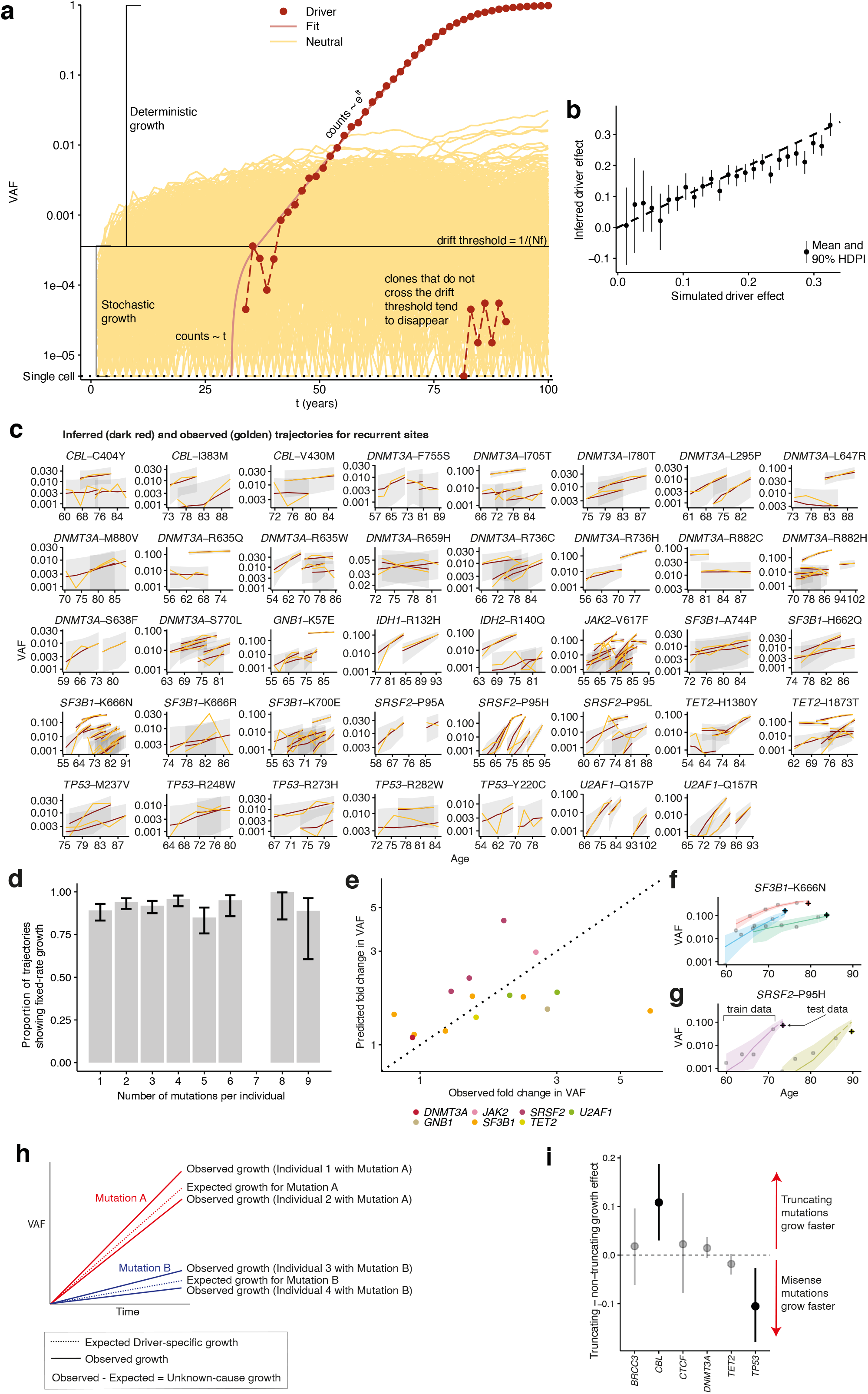
Modelling CH dynamics in older age using time-series VAF data (previous page). **a**, Representation of a Wright-Fisher simulation, showing two phases of clonal growth. The likelihood of a clone transitioning from stochastic to deterministic growth is inversely proportional to the product of its fitness (f) and the total number of stem cells (N). Clones with no fitness advantage (depicted in yellow) are unlikely to exceed their drift thresholds and tend to disappear or remain undetectable. Fitter clones (depicted in red) are more likely to reach deterministic growth. **b**, Association between the driver mutation effect used in the Wright-Fisher simulations and the driver effect inferred using our model (R^2^ = 0.92). **c,** Comparison of observed (golden) and inferred (red) trajectories for all recurrently mutated sites. Grey bands represent 95% highest posterior density intervals. **d**, Relationship between the number of mutations co-occurring within an individual and the proportion of clones growing at a fixed rate over time. **e**, Association between VAF predicted by our model, and VAF observed in additional prospectively-collected samples from 11 individuals with 15 CH driver mutations, not used to infer clonal growth rate in our model. The dotted line is along the diagonal, depicting theoretical perfect agreement between predicted and observed VAF. **f,g**, Example trajectories of clones with *SF3B1*-K666N (f) and *SRSF2*-P95H (g) mutations. Points represent VAFs used in our model to fit the growth curve (train), and crosses represent prospectively tested VAFs used (test), showing good agreement between predicted and observed VAFs. **h**, Illustration of the determinants of growth in our model. Each mutant gene and/or site drives an expected rate of clonal growth. In this example, Mutation A is expected to drive faster growth than Mutation B. The growth rates of different clones bearing the same mutation, either in different individuals or in distinct clones within the same individual, can differ. Some grow faster than expected from the identity of the driver mutation (eg. Individual 1 with Mutation A), and some grow slower (eg. Individual 2 with Mutation A). The residual term in our model, the difference between observed and expected growth rate, is referred to as “unknown-cause growth”. **i,** Comparison of growth rate associated with truncating vs non-truncating mutations in genes with both driver types. Points above the dashed line show faster growth of truncating mutations, and points below show faster non-truncating mutations.

**Extended Data Fig. 5:**
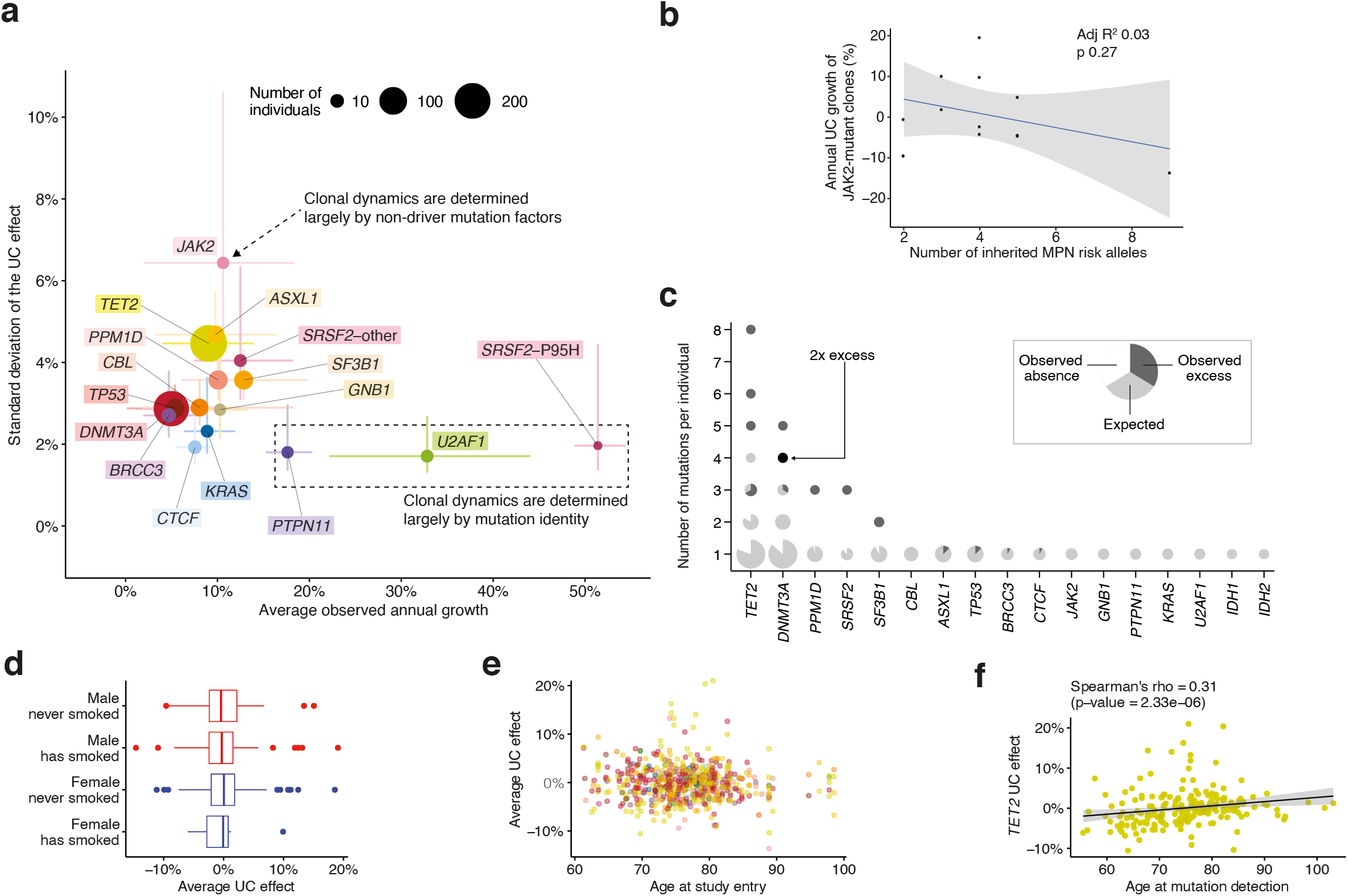
Differences in growth rate between individuals/clones with the same driver. **a**, For each gene, we contrast the mean annual growth rate among individuals/clones bearing a mutation in that gene, with the spread in this rate (defined here as the standard deviation of the unknown-cause (UC) growth). Circles represent point estimates, with circle size indicating the number of clones bearing a mutation in that gene, and lines representing the 90% confidence interval (CI). For the standard deviation, the 90% CI was calculated assuming that 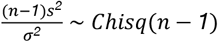, with *n* being the sample size, *s* the standard deviation estimate and *σ*^2^ the true population variance. *SRSF2*-P95H mutations are plotted separately to other SRSF2 mutations, as they are associated with significantly different growth dynamics. **b**, Relationship between number of inherited MPN risk alleles and *JAK2*-mutant clonal growth rate. **c**, The number of mutations per individual in each gene is plotted. Each data-point is a pie-chart, the size of which reflects the number of individuals. For each gene, given the observed mutation prevalence in our cohort, the pie is fully light grey if the number of individuals we observed with a specific number of mutations is the same as the number of individuals we expected by chance. The presence of a white segment indicates that we found fewer individuals with that number of mutations, compared to expected. The presence of a dark grey segment indicates that we found an excess of individuals with that number of mutations. We estimate the expected number of mutations in each gene in each individual through Monte Carlo estimation; assuming the prevalence of mutations in the cohort is uniform for each gene across individuals, we simulate 1,000 scenarios where we randomly distribute these mutations given the number of mutations in each individual. **d,** Association between sex and smoking history and the average UC effect for each individual (n.s.). **e,** Association between age at study entry and the average UC effect for each individual (n.s.). **f,** Association between age at mutation detection and UC effect for each *TET2*-mutant clone (Spearman’s rho = 0.31; p-value=2.33*10^−6^).

**Extended Data Fig. 6:**
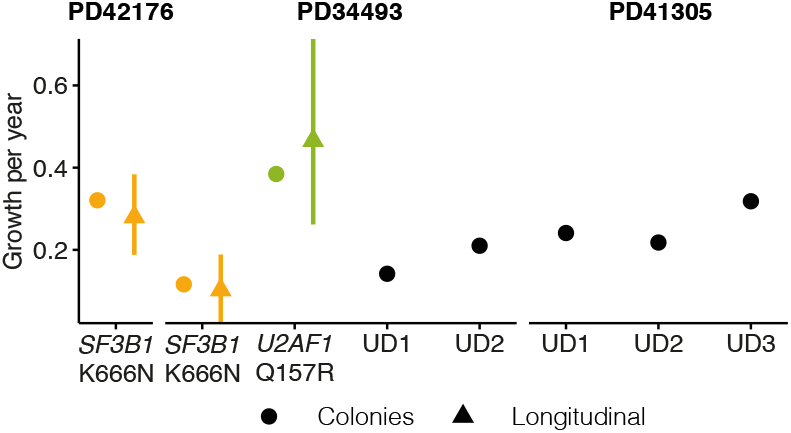
Lifelong growth in phylogenetic trees. Comparison between annual growth derived from phylogenies and growth observed in longitudinal data. For the phylogenies this was obtained by fitting an exponential growth curve to the entire phylodynamic trajectory.

**Extended Data Fig. 7:**
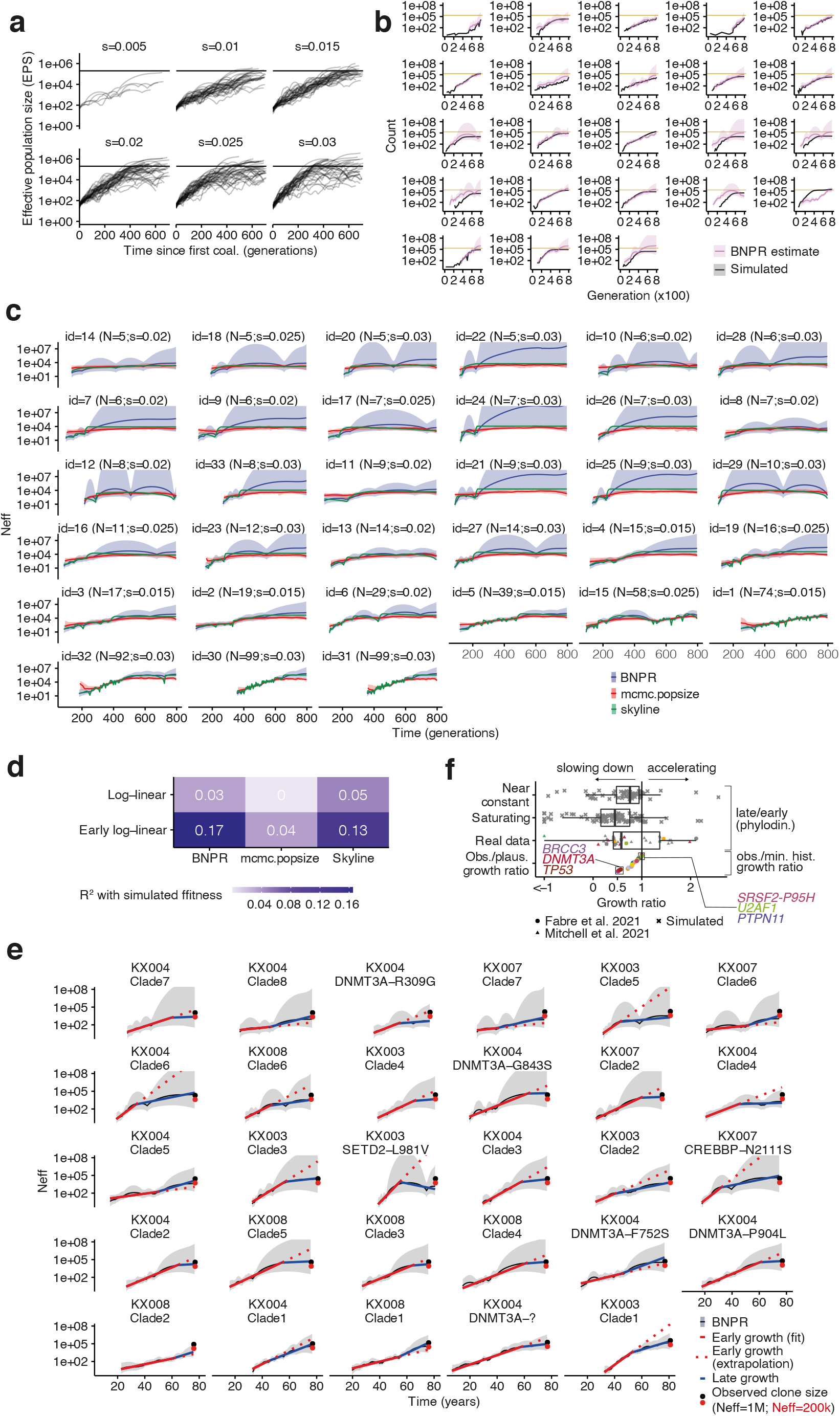
Examples and consistency of clonal deceleration from simulations and real data. **a**, Simulated BNPR trajectories from Wright-Fisher simulations with a fixed population size across 800 generations for a range of fitness effects (0.005, 0.010, 0.015, 0.020, 0.025, 0.030). **b**, Comparison between Wright-Fisher simulations (grey) and BNPR estimates from phylogenies obtained from these simulations (pink). The horizontal golden line in each plot represents the HSC population carrying capacity (200,000). **c**, Representation of effective population size (Neff) trajectories using three distinct methods (BNPR, mcmc.popsize and skyline; details in the Supplementary Methods) for their estimation across a range of clade sizes and fitness effects. **d**, Quantification of the association between true fitness values and inferred fitness values for three distinct methods of Neff estimation. **e**, Schematic representation of all trajectories from Mitchell et al. and how extrapolating from the initial growth rate leads to the overestimation of the observed clone size (here the observed clone size is obtained by scaling the proportion of tips in a clade by a total Neff of either 200,000 or 1,000,000 HSC x yr). **f**, Quantification of the deceleration effect from real data and simulations.

**Extended Data Fig. 8:**
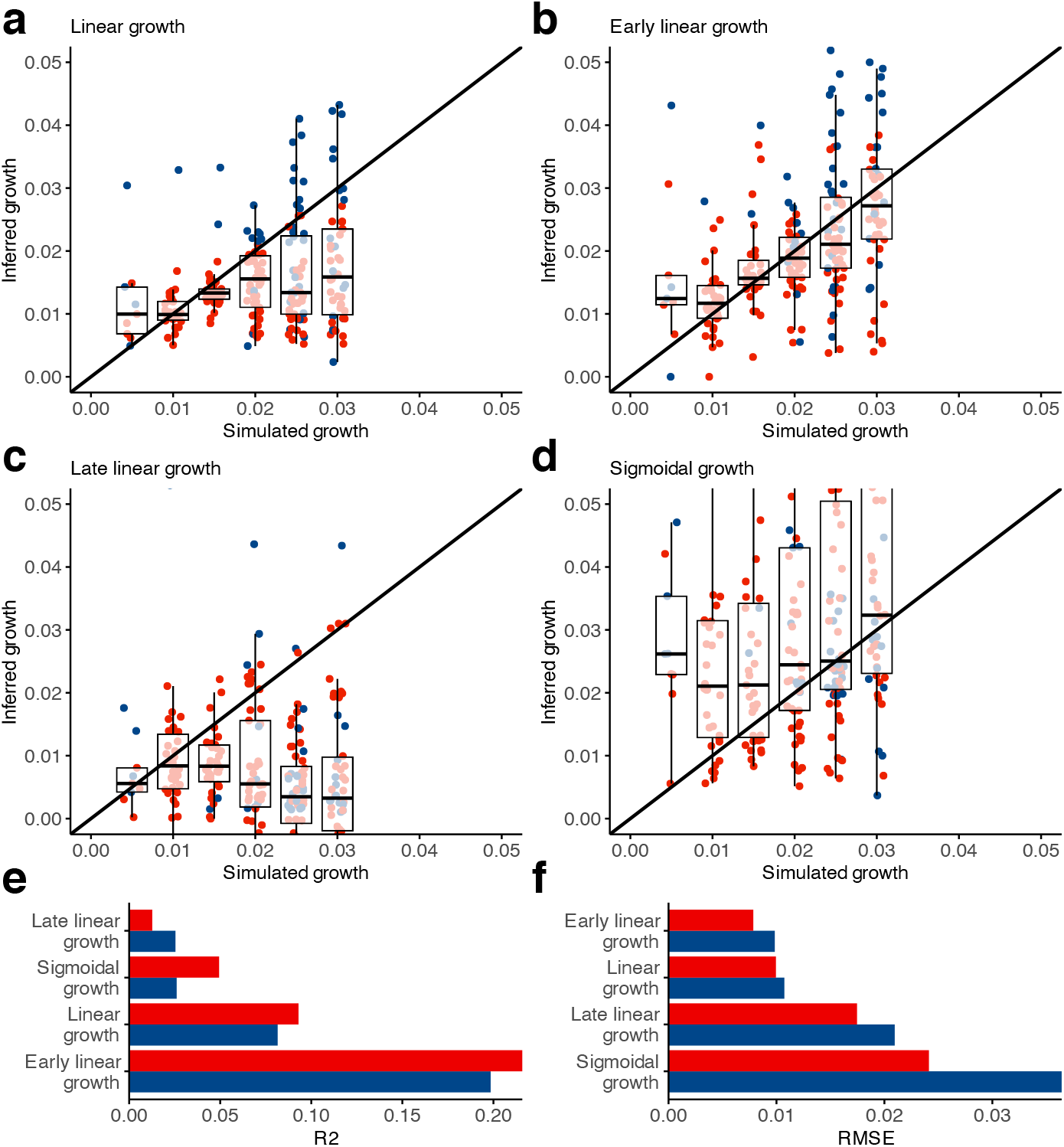
Estimation of the true clone fitness from phylodynamic estimation. Three fits were tested to estimate the true clone fitness from phylodynamic estimation of the population size and these estimates were plotted as a function of the true fitness size (0.005, 0.010, 0.015, 0.020, 0.025 or 0.030). **a**, A log-linear fit; **b-c**, A biphasic fit that estimates an early and a late growth rate and a change-point between both and **d**, a sigmoidal fit. **e**, Coefficient of correlation (R2) for all four inferred coefficients. **f**, Root mean squared error (RMSE) for all four inferred coefficients. In this figure red represents “low variance trajectories” (the average estimated variance for the logarithm of the trajectory is under 5) and blue represents “all trajectories”.

**Extended Data Fig. 9:**
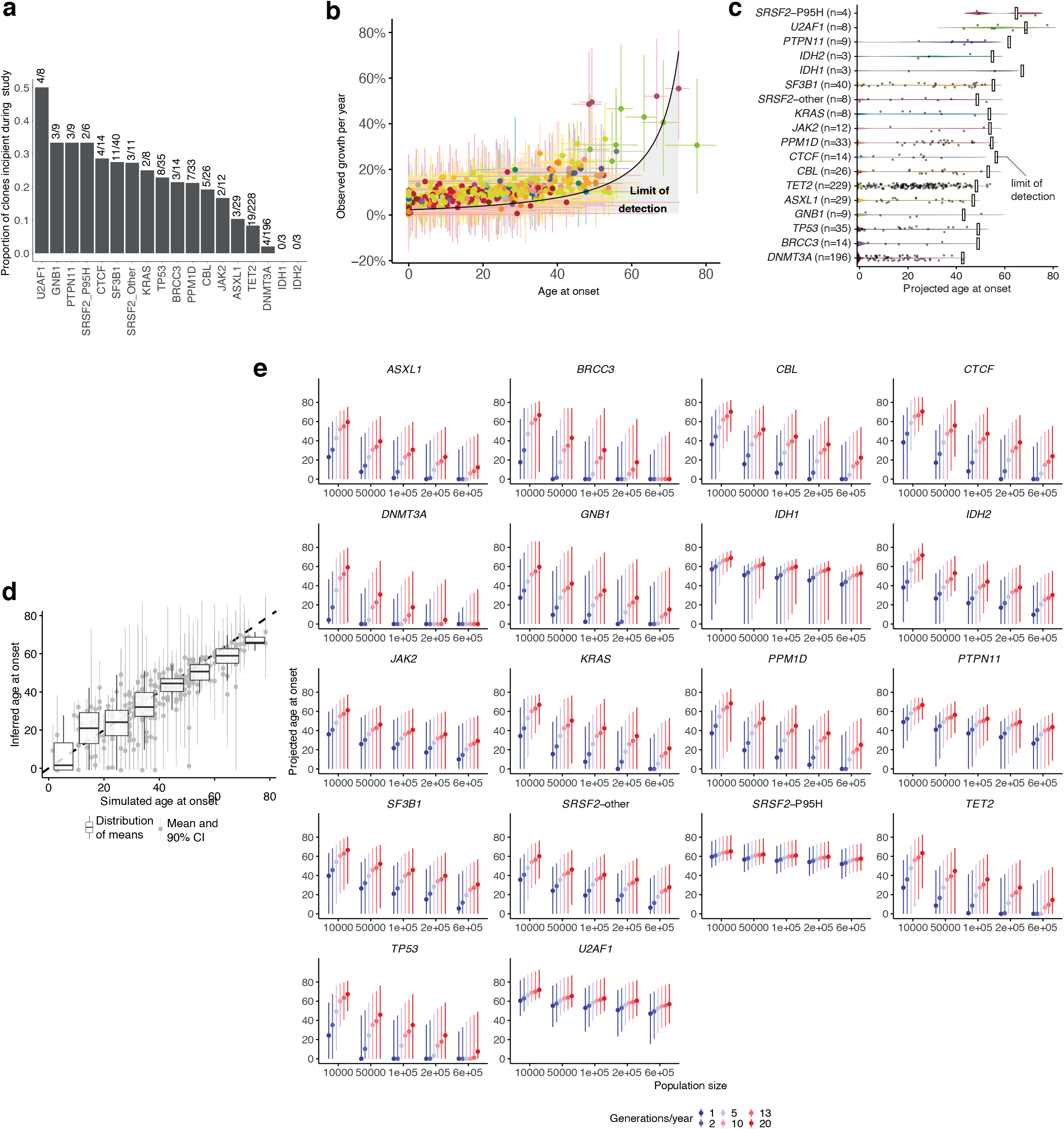
Age at clone detection and onset. **a**, Proportion of clones driven by different driver mutations that were incipient on-study, ie. undetectable at time-point 1 and detectable by the end-of-study. Absolute numbers are given above each bar. **b**, Relationship between age at onset and observed annual growth rate, with 90% highest posterior density intervals (HPDI). The black line and grey shaded area represent the theoretical limit of detection at 80 years of age. **c**, Violin plot showing the distribution of projected ages at onset for all clones, assuming stable lifelong growth at the same fixed rate we observed during older age. **d**, Association between the age at which clones appeared in the simulations and the age at clone foundation inferred using our time-series data (R^2^ = 0.75). Boxplots show that, while these estimates may have high variance, the distribution of expected values is close to the true value. **e**, Sensitivity analysis depicting the median (dot) and the 95% confidence interval of the ages at onset for each gene when considering different population sizes (10e3, 50e3, 100e3, 200e3 and 600e3) and numbers of generations per year (1, 2, 5, 10, 13, 20).

**Extended Data Fig. 10:**
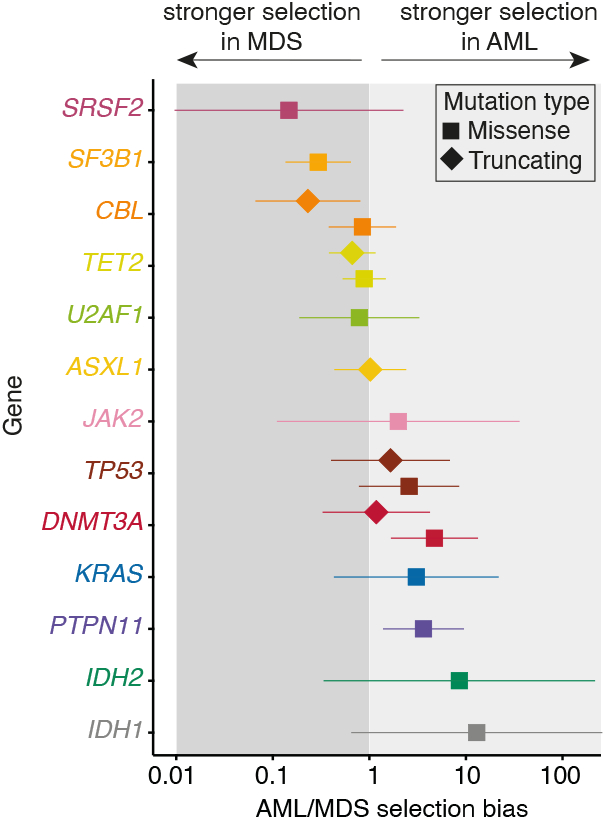
Selection in myeloid malignancies. **a**, Ratio between AML dN/dS and MDS dN/dS for different genes and mutation types (missense, truncating). If this ratio is >1 there is a bias towards AML, if it is <1 there is a bias towards MDS. Confidence intervals for the ratios were calculated under the assumption that dN/dS estimates are normally distributed.

## Supplementary Methods

### Assessing the predictive performance of clonal growth predictions

Using an additional time-point (phase 6) available for 11 individuals with mutations in *CBL* (c.2434+1G>A), *DNMT3A* (P385fs, R882H, W330X), *GNB1* (K57E), *JAK2* (V617F), *PPM1D* (Q524X), *SF3B1* (K666N, K700E, R625L), *SRSF2* (P95H, P95L), *TET2* (Q1542X) and *U2AF1* (Q157P, Q157R). Using the model described in the “Hierarchical modelling of clone trajectories through time” section of the Methods and conditioning on the previous timepoints, we predict the additional time-point and assess the predictive performance through the mean absolute error (MAE) to the true VAF value.

### Validating the dynamic coefficient and age at onset inference with Wright-Fisher simulations

We use Wright-Fisher simulations^1–3^ with a fixed population of 200,000 cells and 50 possible drivers, a range of fitness advantages (*0.001* − *0.030*) and a range of mutation rates (*1.0* ∗ *10* ^*−10*^ *− 4.0* ∗ *10* ^−*9*^). These ranges were estimated to cover the values inferred and mentioned in considering that one should expect there to be approximately 13 generations of HSC per year and a population size of 200,000 HSC ^4^.

To simulate the conditions under which the experimental data was obtained, we fit Gamma distributions to the observed coverage and observed age at first time-point truncated at the minimum and maximum values for each. For each simulation we sample from these distributions the first timepoint, a random number of subsequent timepoints (between 2 and 4) from a uniform distribution and the coverage for each driver at each timepoint. We simulate the sequencing process as drawing samples from a beta-binomial distribution parameterized similarly to the one described in the “Hierarchical modelling of clone trajectories through time” section of the Methods, where the probability is the proportion of cells from a specific clone present at a given time-point. More concretely, 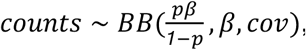, where *p* is the allele frequency of a mutation, *β* is the technical overdispersion parameter and *cov* is the coverage which is sampled from the coverage distribution as inferred from our data.

To infer coefficients under this setting we converted generations to years (13 generations per year) and used the framework described in the previous sections to infer these coefficients. Since the nature of these mutations does not consider different levels of genetic resolution, we had to modify the driver coefficient to 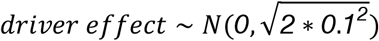so that the distribution from which this coefficient is being drawn has the one we consider for the driver effect considering a gene, domain and site effect. The observed coefficients are converted to year as *coefficients = (1 + fitness)*^*g*^ − *1*, where *g* is the number of generations per year, and we assess the fit between inferred and observed coefficients considering these values. We additionally calculate the age at clone foundation for the inferred coefficients and, using these simulations which allow us to know the true age at clone foundation, we assess the fit between inferred and observed ages at clone foundation.

To better understand the impact that population size and generation times have on these simulations, we conduct the same analysis considering two additional scenarios: a population size of 100,000 HSC and 5 generations per year, and a population size of 50,000 HSC and 1 generation per year.

Finally, we also calculate the age at onset as specified in the “Determining the expected age at beginning of clone onset”. To do this, we assume that these clones follow a Wright-Fisher process, where growth can be separated into two distinct phases which depend on the size of the clone - a stochastic phase, where the clone is too small and during which growth happens linearly, and a deterministic phase, during which growth is approximately exponential (Extended Data Fig. 4a). According to this growth regime, the age at onset can be calculated as 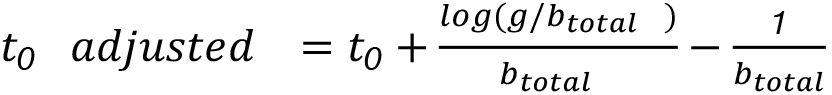, where *t*_*0*_ is the age at onset if the clone grew exponentially (as opposed to following a Wright-Fisher process), 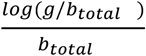 is the time at which the clone started to grow deterministically and 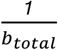 is the expected time the clone spends following a stochastic growth regime. We assess the validity of this approach by calculating the coefficient of correlation between inferred and true ages at onset from the simulations.

### Validating annual growth rate inferences from single-cell phylogenies with Wright-Fisher simulations

We use Wright-Fisher simulations ^2,3^ with 50 possible drivers and test a range of different fitness advantages ([0.005,0.010,0.015,0.020,0.025,0.030]) over 800 generations at a fixed population size of 200,000 HSC. For each fitness effect we define a driver mutation rate ([*200* ∗ *10* ^−*9*^,*50* ∗ *10* ^−*9*^,*20* ∗ *10* ^−*9*^,*15* ∗ *10* ^−*9*^,*8* ∗ *10* ^−*9*^,*5* ∗ *10* ^−*9*^], respectively) that guarantees that at least a few simulations lead to clones which expand to sufficient sizes and avoid many competing expansions and keep the passenger mutation rate constant (*2* ∗ *10* ^−*5*^). For each simulation we infer phylogenetic trees by sampling 100 representative clones from our population and using a neighbour-joining algorithm based on mutation presence. The representative sampling is done by defining for each clone a probability of being sampled that is equivalent to its proportion in the population. We then detect the clades that contain drivers, isolate them and infer their effective population size (Neff) trajectory using BNPR ^5,6^.

We fit different models to the inferred Neff trajectories, namely:

1. A log-linear fit (assumes exponential growth);
2. A scaled and shifted sigmoidal fit (assumes that growth saturates based on the Neff trajectory);
3. A shifted sigmoidal fit (assumes that growth saturates at 1 and that the most recent Neff estimate corresponds to the proportion of tips in the clade);
4. A biphasic log-linear fit (assumes that growth is exponential and has two distinct coefficients corresponding to early and late growth; the boundary between early and late growth - otherwise referred to as the changepoint between both - is also fitted with the other parameters and is constrained to lie in the central part of the trajectory: for the time ) over which the clone expands, the changepoint cannot be inferior to *min*(*t*) + *0.25* ∗ *range*(*t*) nor superior to *max*(*t*) − *0.25* ∗ *range*(*t*), where *range*(*t*) = *max*(*t*) − *min*(*t*). This constraint prevents fits that are too close to the clonal inception or to the clone at later stages).

We compare these models by assessing how closely they are able to recapitulate the original fitness in the simulations. To do so, we calculate their coefficient of determination and root mean squared error. We also visually assess how similar these trajectories are to the true driver trajectories as reconstructed from simulations - to match clones from a Wright-Fisher simulation to an expansion in a phylogenetic tree we assign each clone from the Wright-Fisher simulation to its nearest clone in a phylogenetic tree using the Hamming distance between the mutations in each clone.

We additionally estimate the effective population size using two other methods for validation - mcmc.popsize and skyline from the ape package ^7^ in R. This allows us to confirm our observations that stem from phylodynamic estimations and that concern, mostly, a prevalent effect of clonal deceleration which is detailed in the main text and in the following section.

### Detecting deceleration in single-cell phylogenies and longitudinal data

We infer the presence of deceleration in both single-cell phylogenies and longitudinal data. To do this, we use two distinct methods: calculating the ratio between the expected and observed VAF and calculating deceleration using growth rates.

For the first method - calculating the ratio between expected and observed VAF - we use the value for the early growth from the changepoint log-linear fit described in “Validating annual growth rate inferences from single-cell phylogenies with Wright-Fisher simulations” and extrapolate the Neff to the age at sampling. By doing so we get the expected clone fraction if growth had not changed during the Neff trajectory. We also calculated the observed clone fraction as the fraction of tips in the clade. To get the expected clone fraction from Neff we divide Neff by the inferred population size in Lee-Six et. al (200,000 HSC) ^8^. We then calculate the ratio between the expected and observed clone size - if this ratio is close to 1 this implies little to no changes in dynamics, whereas a ratio above 1 implies deceleration and a ratio below 1 implies acceleration.

For the second method - calculating deceleration using growth rates - we define two distinct quantities for both single-cell phylogenies/longitudinal data - expected/observed growth, corresponding to the growth rate of each clone during observation at old age, and early/minimal historical growth, corresponding to the growth rate of each clone at an earlier stage of clonal dynamics - and calculate the ratio between them.

As such, for phylogenies we first calculate the Neff trajectory for each clade using BNPR ^33^. Next, and using their Neff trajectory, we calculate their expected growth rate by assuming a sigmoidal growth. We additionally assume that the final Neff (Neff at sampling) estimate corresponds to the fraction of tips in the clade and we scale our data accordingly such that 1 corresponds to the maximum Neff and the fraction of tips in the clade corresponds to Neff at sampling. Thirdly and using the changepoint log-linear fit described in “Validating annual growth rate inferences from single-cell phylogenies with Wright-Fisher simulations” we derive the value for early growth. Finally, as a measure of deceleration, we calculate the ratio between expected and early growth - a value close to 1 for this ratio implies an absence of deceleration whereas smaller values imply deceleration.

For the longitudinal data we use the observed growth for each clone as described in “Hierarchical modelling of clone trajectories through time”. Next, we calculate the (minimal) historical growth as the growth that excludes all posterior samples that would lead to age at onset estimates exceeding lifetime (ages at onset for clones below −1, a heuristic value chosen to represent developmental onset of clones). Finally and as a measure of deceleration, we calculate the ratio between observed and historical growth. The interpretation for this ratio is similar to that defined in the previous paragraph for phylogenetic data - a value of 1 implies an absence of detectable deceleration, whereas smaller values represent the minimal amount of deceleration. This method has, however a caveat - due to the nature of this calculation (excluding posterior samples which are too slow to provide solutions within lifetime), values above 1 (indicating acceleration) are technically impossible.

## Supplementary Notes

### Supplementary Note 1 - Determining the effect of repeated sampling on the theoretical limit of detection

Across this work we sequence individuals a median of three times across their lifetime. We define a detection threshold of 0.5% VAF as the minimum clone size for detection on individual timepoints, but the repeated sampling leads to 0.5% VAF being an overestimation of the actual limit of detection (LOD) - the size at which clones become detectable.

To show this, we simulate the repeated sampling of variants existing at a true clone proportion between 0 and 2%. We use this proportion *p* as the probability parameter in a beta binomial distribution, the overdispersion *β* calculated using technical replicates as the overdispersion in the same beta binomial distribution and a coverage of 1000. Having fully parameterized this distribution 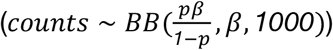 we sample counts from it between 1 to 5 times. For each combination of clone size and number of samples we perform 1,000 realisations and calculate the number of detected clones at a threshold of 0.5%. This allows us to assess the fraction of clones with a specific size which are detected if we sample them multiple times - in other words, are able to assess the detection rate for different clone sizes and different numbers of samples.

With this, we show that, at a threshold of 0.5% and sampling only once, we detect 14.8% of all clones existing at 0.5% (Supplementary Notes Fig. 1). However, repeating this sampling 3 and 5 times leads to the detection of approximately 37.7% and 54.3% of all clones existing at 0.5%, respectively. As such, under regular conditions - a single sample - we would detect 13.5% of all clones present at 0.5% with a detection threshold of 0.5%. The question we should now ask is: what is the smallest possible clone size we detect at the same rate of detection - 13.5% - if we increase the number of samples? Using the same set of simulations, we can calculate the likely minimal size of the detected clones, summarised in Supplementary Notes Table 1, with clones as small as 0.21% and 0.14% being detected with 3 and 5 samples, respectively, using the same detection rate. As such, when considering the theoretical LOD used in Figure 4k, we avoided using 0.5% which, as we show, would be at least twice as high as the theoretical LOD obtained from simulations.

**Supplementary Notes Fig. 1.**
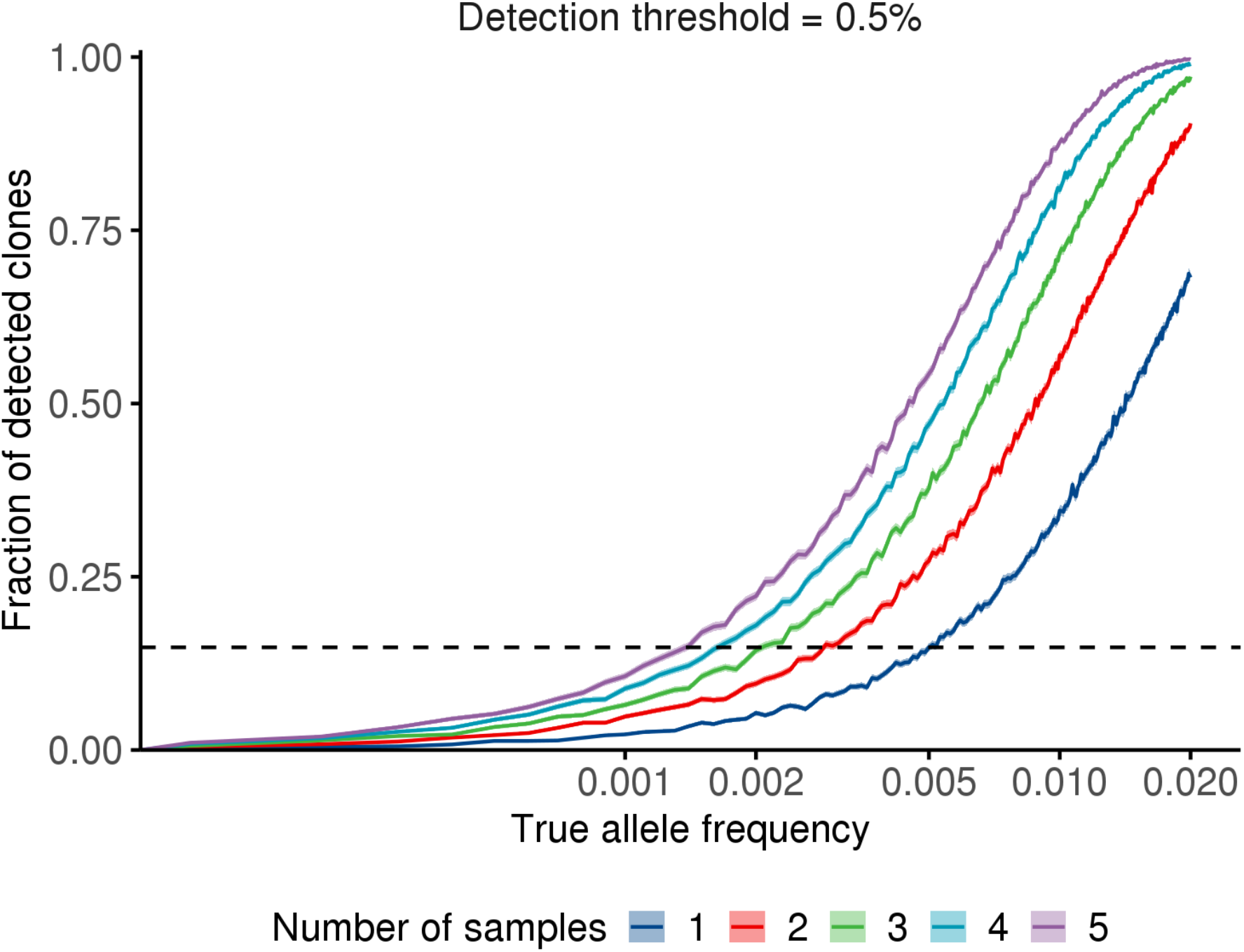
Fraction of detected clones upon repeated samples/timepoints at a detection threshold of 0.5%.

**Supplementary Notes Table 1.** The minimal size of detected clones using a 0.5% threshold and assuming that we are interested in detecting the same fraction of clones we would detect with a single sample at a detection threshold of 0.5%.

| Number of samples | Minimal size of detected clones at 15.08% |
| --- | --- |
| 1 | 0.50% |
| 2 | 0.30% |
| 3 | 0.21% |
| 4 | 0.16% |
| 5 | 0.14% |

